# Unravelling the effects of heterogeneity in space use on estimates of connectivity and population size: Insights from spatial capture-recapture modelling

**DOI:** 10.1101/2024.03.04.583266

**Authors:** Maëlis Kervellec, O. Gimenez, C. Vanpé, P.-Y. Quenette, J. Sentilles, S. Palazón, I. Afonso Jordana, R. Jato, M. M. Elósegui Irurtia, C. Milleret

## Abstract

Spatial Capture-Recapture (SCR) models using least-cost path distance offer an unified framework to estimate landscape connectivity and population size from individual detection data, while accounting for individual and spatial heterogeneity in space use. In our case study on the Pyrenean brown bear population, two “outliers” individuals were detected more often over very large spatial extent compared to other individuals. The integration of such individuals made SCR modelling challenging, especially since it remains unclear how unmodelled heterogeneity in space use may bias connectivity and population size estimates. To address this gap, we used simulations reflecting the Pyrenean brown bear population, with two groups of individuals differing in their space use due to individual characteristics but also in their spatial responses to landscape structure. We compared six SCR model formulations that varied in whether individual and spatial heterogeneity in space use were (1) ignored, (2) explicitly modelled, or (3) handled by removing outliers. We then applied the same models to the empirical bear dataset.

By combining simulations and sensitivity analyses, we highlight the challenges of choosing the appropriate modelling approach when multiple sources of heterogeneity in space use occur simultaneously. Our results show that the treatment of heterogeneity in space use should match the research objective: removing outliers supports accurate population size estimation, while explicitly modelling heterogeneity is essential for reliable connectivity assessment. Overall, we provide a practical framework for identifying and addressing heterogeneity in SCR applications, to guide practitioners in deciding when additional model complexity is warranted.

## 1. Introduction

Animals move through the landscape according to intrinsic (e.g., locomotion abilities, need to feed or to find mates) and extrinsic factors (e.g., resource availability, risk to encounter predators) (Bell, 1990). This means that animals do not move randomly through the landscape, but constantly have to adjust their movement to the distribution of opportunities and pressures. Identifying the spatial features affecting animal movement can be done by quantifying connectivity. Connectivity aims at describing “the degree to which the landscape facilitates or impedes movement among resource patches” (Taylor et al., 1993). Population-level connectivity can be estimated using a map called a resistance surface that quantifies the cost of moving through each pixel of the landscape. This surface is estimated from spatial environmental variables and information on the cost of movement or space use derived from expert opinion or empirical animal movement data (Dutta et al., 2022; Zeller et al., 2012). In landscape management, resistance surfaces are used to map corridors, detect barriers or predict population range shift due to climate change (Dutta et al., 2022). However, connectivity does not only have a spatial dimension, but also varies with time (Zeller et al., 2019), or with individual characteristics: e.g. life stages (Thorsen et al., 2022), or sex (García-Sánchez et al., 2022; Zeller et al., 2023). Despite the importance of intraspecific variation, group specific connectivity is rarely considered (Garcia-Sanchez et al., 2022; Zeller et al. 2023).

Spatial capture-recapture (SCR) models, which have become standard in population ecology (Efford, 2004; Tourani, 2021), are hierarchical models that use spatially explicit individual detection histories to estimate both population size and distribution. A core assumption of SCR models is that detection probability declines with distance from an individual”s activity centre (Royle et al., 2014). In other words, the detection function is designed to reflect individual space use, as it shapes where and how often an individual is detected. In the context of SCR models, we define space use as the spatial distribution of detection probability, with baseline detection probability set to 1 (*p*_0_ = 1), and conditional on the estimated location of the activity centre. In its most commonly used form, a SCR model assumes a circular space use resulting in a symmetrical and stationary (invariant across space) space use extent. However, extensions of SCR models allow the estimation of nonstationary, asymmetrical individual space use extent caused by spatial heterogeneity in the landscape (Royle et al., 2013). In these models, distance in the detection function is not Euclidean but based on the least-cost path (LCP) distance, with landscape resistance directly inferred from the data. As a result, SCR models with LCP distance provide a unified framework for jointly estimating both population size and the effects of landscape resistance on space use (Royle et al., 2013; Sutherland et al., 2015), thereby quantifying population-level landscape connectivity. This approach has been applied to several species, including American mink (*Neogale vison*, Fuller et al., 2016), black bear (*Ursus americanus*, Morin et al., 2017), jaguar *(Panthera onca*, Tobler et al., 2018), snow leopard (*Panthera unica*, Pal et al., 2021), and brown bear (*Ursus arctos*, Kervellec et al., 2023). However, these studies focused on population-level connectivity and did not account for individual heterogeneity in space use, particularly in the resistance parameter that scales the spatial heterogeneity in space use.

Individual heterogeneity refers to variability in responses among individuals within a population, which may be linked to measurable or non-measurable individual characteristics (Gimenez et al., 2018). In non-spatial capture-recapture models, it is well established that accounting for individual heterogeneity in detection probability is crucial to prevent negative bias in population size estimates (Gimenez et al., 2018). In SCR models, heterogeneity in detection probability can arise from individual variation in the baseline detection probability as well as from individual and spatial heterogeneity in space use. Our focus here is on the latter. Non-Euclidean SCR models typically describe heterogeneity in space use with two parameters: (1) the scale parameter *σ*, which defines the extent to which an individual can be detected from its activity centre and is related to home range size (Royle et al., 2014), and (2) the resistance parameter *α*_2_, which quantifies the influence of spatial covariates on space use across the landscape (i.e. spatial heterogeneity). Therefore, the scale parameter *σ* can vary among individuals to reflect individual variation in space use while the resistance parameter *α*_2_ quantifies how individuals adjust their space use to spatial heterogeneity. Not all individuals may respond similarly to landscape structure, leaving the possibility for individual-specific *α*_2_ parameters. Accounting for individual heterogeneity in the scale parameter is important to prevent negative bias in population size estimates (Gardner et al., 2010; Sollmann et al., 2011). SCR models are generally robust to unmodelled spatial heterogeneity in space use (Royle et al., 2016; Theng et al., 2022) but strongly structured landscapes can nevertheless cause an underestimation of population size (Sutherland et al., 2015). In addition, unmodelled individual heterogeneity in the response of space use to landscape structure remains unknown. Although previous studies have shown that unmodelled heterogeneity in space use, whether arising from individual traits or from the location of individuals in the landscape, can bias population size estimates (Gardner et al., 2010; Sutherland et al., 2015), much less is known about the combined effects of individual and spatial heterogeneity on estimates of both population size and connectivity.

The motivation for this study stems from our work on the Pyrenean brown bear population (Kervellec et al., 2023; Sanz-Pérez et al., 2025). Nearly extinct in 1995, this population is now recolonising human-dominated landscapes (Kervellec et al., 2023; Sanz-Pérez et al., 2025). In this context, robust estimates of population size and a clear understanding of how landscape structure affects connectivity are critical. In a previous study, Kervellec et al. (2023) identified that two individuals were detected far more frequently and had a much larger apparent space use than the rest of the population. The common practice in this case is to remove these “outliers” individuals from the analyses, in order to avoid biased population size estimates due to unmodelled heterogeneity in the space use component of the detection probability (Kendall et al., 2019; Schmidt et al., 2022). However, such wide-ranging individuals may in fact play a key role when the goal of the study is to quantify population connectivity.

In this study, we investigate how heterogeneity in space use affects (1) population size estimate and (2) connectivity estimates, using non-Euclidean SCR models. For each objective, we built six different models to evaluate the consequences of (a) ignoring individual or/and spatial heterogeneity in space use or (b) removing the individuals with large space use (“outliers”). We first conducted a simulation study inspired by the Pyrenean brown bear population, with two groups of individuals differing in both space use extent and response to landscape structure. We then applied the same models to the empirical Pyrenean brown bear data. We expected that (1a) population size would be underestimated when individual heterogeneity in space use is not accounting for (Gimenez et al., 2018; Sutherland et al., 2015), (1b) removing outliers would yield unbiased population size estimates, and (2) removing outliers and not accounting for individual heterogeneity would undermine estimates of space use and connectivity.

## 2. Materials and Methods

### 2.1 Spatial Capture Recapture models to estimate connectivity

Spatial capture-recapture (SCR) models are hierarchical models that estimate the location of each individual activity centre denoted s_i_ in the spatial domain S according to a spatial point process. This process can be homogeneous, in this case s_i_ ∼ Uniform(S) or can be driven by spatial variables. For inhomogeneous point process, an intensity function λ(s) = e^βX(s)^ describes the placement of activity centres, according to the vector of regression coefficients β and the vector of spatial covariate X(s) evaluated at cell s (Zhang et al., 2023). SCR models combine an ecological sub-model describing the distribution of the population”s activity centres and observational sub-model. The observation sub-model is based on the detection and non-detection of individuals through an array of traps and account for the imperfect detection of individuals by explicitly estimating detection probability (Efford, 2004; Royle et al., 2014). Therefore, the detection history corresponds to detecting or not the individual i at trap j at the sampling occasion k in session t, and is noted y_ijkt_:

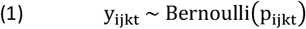

Where p_ijkt_ is the detection probability usually modelled by a half-normal function:

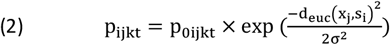

In this relation, the detection probability decreases with the Euclidean distance, noted d_euc_;x_j_, s_i_), between the trap x_j_ and the activity centre s_i_. The baseline detection probability p_0ijkt_ is the probability of detecting an individual if its activity centre is located exactly on the trap. The scale parameter σ provides the extent to which an individual can be detected and is correlated with space use extent. The larger the σ, the further from its activity centre an individual can be detected. These parameters can be constant or varying according to individual covariates (e.g., sex) to account for individual heterogeneity in the detection process.

By using the Euclidean distance, we assume that space use is symmetric and stationary (Royle et al., 2014), meaning that the presence of barriers or resources do not affect movements of individuals across the landscape. We account for the effect of landscape structure by assuming that detection probability is a function of the ecological distance instead of Euclidean distance. The ecological distance can be represented by the least-cost path distance, which is based on a discrete resistance surface, where each pixel, noted ν_g_, is given a cost. The distance between two pixels in the landscape is the succession of m steps, noted ν_1_, ν_2_,…, ν_m_, forming a path. We calculate the cost of all possible paths (ℒ_1_, ℒ_2_,…, ℒ_l_) connecting these two pixels and keep the one with the minimum value:

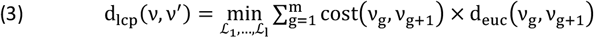

where, cost 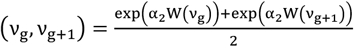

The resistance parameter α_2_is explicitly estimated in the SCR model from the encounter data and W(ν) is the resistance variable considered to build the resistance surface at cell ν. The estimated resistance parameter reflects the extent to which W decreases (α_2_ > 0) or increases (α_2_ < 0) the space use extent of individuals across the landscape (Royle et al., 2013; Sutherland et al., 2015). When α_2_ = 0, the resistance is null and the distance is exactly Euclidean. The resistance parameter therefore quantifies the extent to which spatial characteristics of the landscape influences individuals space use (spatial heterogeneity in space use). Similarly to σ, α_2_ can vary according to individual characteristics according to a function of individual covariates (spatial and individual heterogeneity in space use). Here the least-cost path distance is computed by the Dijkstra algorithm implemented in the *gdistance* R package (version 1.6.4, van Etten, 2012).

A map of the potential landscape connectivity can be derived from these estimates. Given the location of an activity centre, we can compute its space use from equation 2, by setting p_0_ = 1. We then assume that an activity centre is present in each pixel of the landscape and compute the sum of the space use of each individual at each pixel (Morin et al., 2017) to obtain a measure of potential connectivity. Therefore, potential connectivity goes from 0, meaning that not a single individual can reach this pixel, to the total number of pixels, where all individuals can reach this pixel. A realized measure of landscape connectivity is computed by weighted the potential connectivity map by the estimated number of activity centres in each pixel (i.e. realized densities), also called density weighted connectivity (Morin et al., 2017).

### 2.2 Simulations

We conducted a simulation study to assess the consequences of not accounting for individual and spatial heterogeneity in space use on parameter estimation. We considered a population composed of two groups differing in their responses to landscape structure. The study area was a 25 × 25 distance unit (du) square domain, within which we centred an array of 100 traps spaced by a minimum of 2du. We added a 3.5du buffer around traps, following standard recommendations (Royle et al., 2014). To represent the landscape structure, we defined the resistance covariate by a raster of 0.5×0.5du resolution representing landscape fragmentation with the distance to a road (see map Appendix S1).

The simulation study was designed to mimic issues encountered with the Pyrenean brown bear population (Kervellec et al., 2023). We simulated 100 individuals, assuming a homogeneous distribution of their activity centres within the spatial domain. The population was composed of two groups. The first encompassed 97% of the population (n = 97) and was called the “common group”, while the second one gathered the outlier individuals (n=3) characterized by a larger space use and was called the “outlier group”. We assumed that the larger space use of the “outlier” group was linked to individual variation in home range size (σ_Com_ = 7; σ_Out_ = 5), as well as a different response to the road. The common group have smaller space use when the distance to road decreased 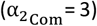 while the outlier group was not affected 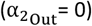. To ensure realistic (i.e. number of detections per individual ≤ 30) detectability rates between the two groups despite their different space use, we also set a higher baseline detection probability for individuals of the common group 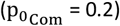 compared to the outlier group 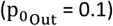.

We simulated 25 detection histories. For each simulation, we fitted six models: four using the full data (denoted MOx) and two with all detections of all outlier individuals removed (denoted Mx). We considered models assuming an effect (MO3, MO4, M4) and no effect (MO1, MO2, M1) of landscape structure on space use. For datasets including outliers (MOx), we sequentially increased the number of model parameters (*p*_0_, *σ, α*_2_) that accounted for individual heterogeneity through a group effect (Table 1).

**Table 1.** Models formulas used for the simulation study and applied on each simulated datasets. It present the model name, the dataset, the baseline detection probability (p_0_), the scale parameter (σ), the resistance parameter (α_2_) when applicable. The empty cells means that the resistance parameter was not included in the model and therefore we used the Euclidean distance instead of the least-cost path distance.

| Model name | dataset | $p_0$ | $\sigma$ | $\alpha_2$ |
| --- | --- | --- | --- | --- |
| MO1 | all | ~1 | ~1 |  |
| MO2 | all | ~group | ~group |  |
| MO3 | all | ~group | ~group | ~1 |
| MO4 | all | ~group | ~group | ~group |
| M1 | outlier removed | ~1 | ~1 |  |
| M4 | outlier removed | ~1 | ~1 | ~1 |

The simulations were performed using the R package *nimble* (version 1.01 de Valpine et al., 2017). We ran two chains of 50,000 iterations each and discarded the first 1,000 iterations as burn-in. For the simulation study, we discarded models that did not converge, based on visual inspection of the trace plots, the Gelman–Rubin statistic 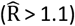 and effective sample size (n_eff_ < 100) (Brooks and Gelman, 1998). We used a data-augmentation approach, with an augmented population of 150 individuals (Royle et al., 2007), and priors are presented in the Appendix S2 (Table S2.A). For the six models, we compared parameter estimates with the true values and recorded whether 95% credible intervals encompassed the truth. We also illustrated differences in connectivity estimates by mapping potential connectivity for models MO3, MO4 and M4 with estimates from the simulated dataset 1. because the 25 simulated datasets were primarily intended to identify trends, we did not compute bias explicitly.

### 2.3 Case study: Brown bears in the Pyrenees

#### 2.3.1 Data

In this case study, we used the spatial capture-recapture dataset of the Pyrenean brown bear population monitored across the Pyrenees mountains in France, Spain and Andorra, presented in Kervellec et al. (2023). In this transboundary study, we collected data from three sampling sessions from 2017 to 2019, in which we defined seven sampling occasions i.e., the months from May to November. We used the data from both the structured and opportunistic monitoring of the population (Vanpé et al., 2022). The structured monitoring consisted of a total of 708 DNA hair snags set on walking transects, around 10% of which are combined with a camera trap (Figure 1). The opportunistic monitoring consisted of 246 traps, defined according to a 25km^2^ grid over the Pyrenees. An opportunistic trap is the centre of each grid cell where at least one depredation of a bear to livestock or beehives occurred in the cell between 2010 and 2020. The depredation events are reported by the owner. In order to get financial compensation, evidences of bears presence and genetic samples (hair or scat) are collected after the depredation (Figure 1). Each sample was genetically analysed following details available in Vanpé et al. (2022), to obtain the identity and sex of the bear detected at a given trap in a given sampling occasion. To account for individual heterogeneity, we divided the population into three groups, females, males and outliers. The maximum distance moved per year, which is the maximum distance between the centroid of the detections and the detections, for Néré was between 39.8 and 66.5km and was between 27.6 and 51.1km for Goiat. The maximum distance moved per year per individual between detections was between 0 (i.e. individuals detected once) and 24.3km for the females (mean = 3.5km) and between 0 and 17.4km for the males (mean = 6.1km) (Figure 2). In average Néré was detected 9 times, Goiat 35.7 times, while males were detected 5.6 times and females 4.1 times per year.

**Figure 1.**
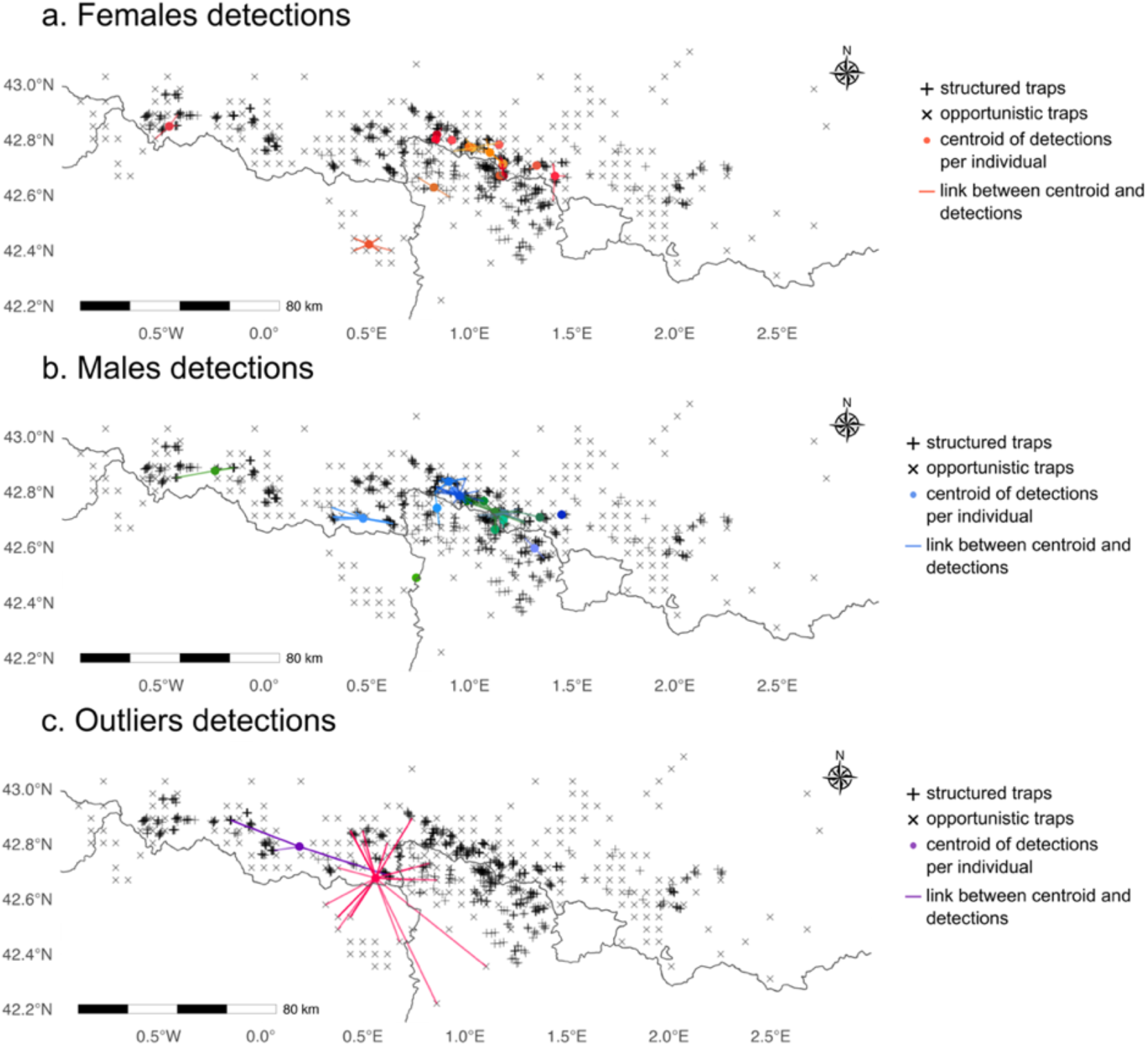
Maps of traps distribution from the structured (+) and the opportunistic (x) monitoring across the Pyrenees mountains (France, Spain, Andorra). Points represents the centroid of detections per individuals and the lines links the centroid to the centroid to the traps where each individual was detected in 2019 for females (a), males (b) and outliers (c).

**Figure 2.**
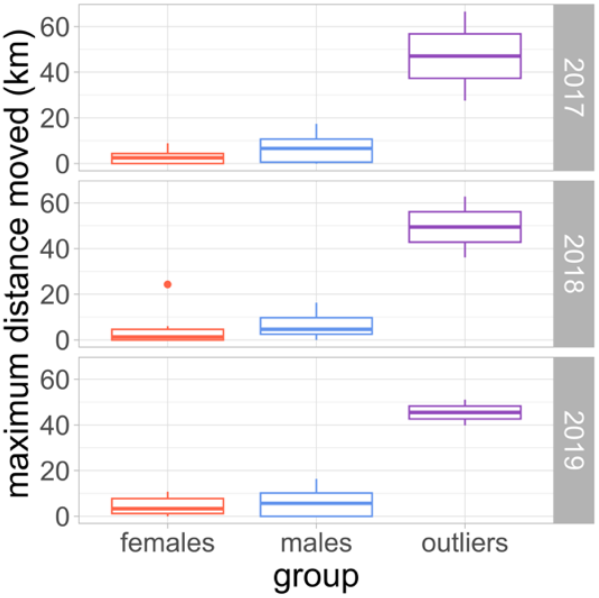
Boxplot of the maximum distance moved in km for each group (females, males and outliers) between 2017 and 2019 for the Pyrenenan brown bear population. This distance is the maximum distance between each detection and the centroïd of its detections for each individual

#### 2.3.2 Models

We applied a SCR model similar to the one used in Kervellec et al. (2023). We assumed that the structured and opportunistic monitoring followed different detection processes such as:

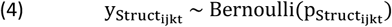

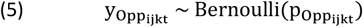

where 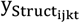 and 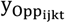 are the encounter histories and 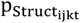 and 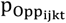 are the detection probabilities for the structured and opportunistic monitoring, respectively. We modelled heterogeneity in the detection probability arising from differences in the sampling protocols or according to brown bear ecology on the baseline detection probability (p_0_) and the scale parameter (σ) using Equation 2.

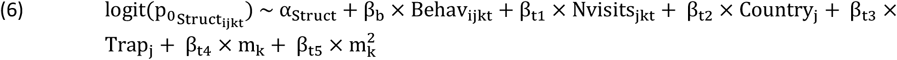

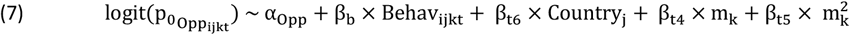

The baseline detection probability (Equation 6) for the structured monitoring was assumed to be a function of a quadratic effect (β_4_, β_5_) of the sampling occasion (m) to account for higher detectability in the summer, the country (Country, β_2_), the number of visits at the trap each month (Nvisits, β_1_) as samples can be collected once or twice each month, and if the hair snag was combined or not with a camera trap (Trap, β_3_). For the opportunistic monitoring (Equation 7), the baseline detection probability was a function of the country (Country, β_6_) and the quadratic effect of the month (m, β_4_, β_5_). Both monitoring accounted for a behavioural effect (Behav, β_b_), meaning that an individual detected once at a trap within a session was more likely to be detected at this trap in the following sampling occasions (Kervellec et al., 2023). We also estimated a different intercept for the structured (α_Struct_) and opportunistic sampling (α_Opp_).

Additionally, the scale parameter was estimated for each sex: σ_i_∼α_s_ + β_s_ × sex_i_, for each group: σ_i_∼α_s_ + β_s_ × group_i_, or was constant (Table 3).

The brown bear densities were assumed to vary according to an inhomogeneous binomial point process:

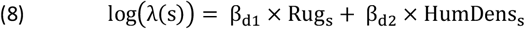

The intensity function (Equation 8) was a function of the linear and additive effect of ruggedness (Rug), defined as average of the absolute values of elevation differences between the focal cell and the eight surrounding cells, and human population density (HumDens) (see Kervellec et al., 2023).

In this Bayesian formulation of the model, we used a data-augmentation approach. The augmented population M has to be large enough to encompass all individuals that were never detected (Royle et al., 2007). Here, we used M = 204 individuals, which is four times the number of detected individuals. The latent state variable z_i_ model the inclusion of an individual from the augmented population to the population with an inclusion probability Ψ:

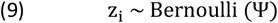

From this equation we can derive the total population size 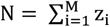, which is the sum of the individuals included in the population.

Spatial heterogeneity in space use was modelled using the LCP distance instead of the Euclidean distance using Equation 3. The resistance surface was based on road density, computed as the length of all roads within each 2.5×2.5km pixel extracted from Open Street Map (see Kervellec et al., 2023). We considered two models with no effect of spatial heterogeneity on space use, but considering individual variation with an effect of sex (MFO1) and group (MFO2) on *σ*. The group is a categorical variable that differentiate between females, males, and outliers. For the other models, we added an effect of spatial heterogeneity in space use assuming that all individuals responded similarly to landscape structure (MFO3) or that response to landscape structure was a function of the group (MFO4). We also added two models where the outliers were removed from the dataset, first modelling only individual heterogeneity in *σ* between males and females with no effect of spatial heterogeneity on space use (MF1) and, second modelling individual and spatial heterogeneity in space use (MF4).

We assumed that we know the group of each detected individual. However, for undetected individuals their group η_i_ is estimated according to a categorical distribution:

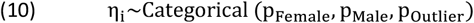

Where *p*_*Female*_, *p*_*Male*_ and *p*_*Outlier*_ are respectively the probability to belong to the group female, male or outlier.

We used the R package *nimbleSCR* (Bischof et al., 2020; Turek et al., 2021). We run three chains of 30,000 iterations each and discarded the first 1,000 iterations as burn-in for each model, and priors are presented in Appendix S3 (Table S3.A). We checked model convergence, based on visual inspection of the trace plots, the Gelman–Rubin statistic 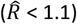 and effective sample size (n_eff_ > 100) (Brooks and Gelman, 1998). We derived potential connectivity maps and density weighted connectivity maps from the distributions of model MFO3 and propagated associated uncertainty. All the analysis were conducted in R (R Core Team, 2023, version 4.3.1)

## 3. Results

### 3.1 Simulation

The simulated data included three outlier individuals that were always detected, while in the “common group” between 42 and 59 individuals were detected out of the 97 present in the population. As expected from the parameter settings, outliers were detected more frequently, with an average of 12.7 detections per individual compared to 2.7 in the common group. All models converged with 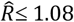 and n.eff > 113.

#### 3.1.1 Population size estimates

Estimates of population size were underestimated when we ignored individual and spatial hetereogenity in space use (Figure 3). When keeping all the individuals in the dataset, the model MO1, that was not accounting for individual and spatial heterogeneity in space use, consistently underestimated population size (95%BCI never cross the true value). Introducing individual heterogeneity in space use (model MO2) led to a mean estimate below the true value in all simulations, with the 95%BCI overlapping the true value for only 0.08% of the simulations. Accounting for spatial heterogeneity in space use (model MO3) resulted in 92% of 95%BCI overlapping the true value. When modelling individual heterogeneity on the resistance parameter (model MO4), 24% of simulations (8/25) were below the true value, and 92% of 95%BCI overlapped the true value (Figure 3). When the three outliers were removed from the dataset, the null model that ignored spatial and individual heterogeneity in space use (model M1) underestimated population size, with 95%BCI overlapping the true value in only 0.08% of the simulations. By contrast, model (M4) that integrated individual and spatial heterogeneity in space use achieved 92% overlap (23/25 simulations).

**Figure 3.**
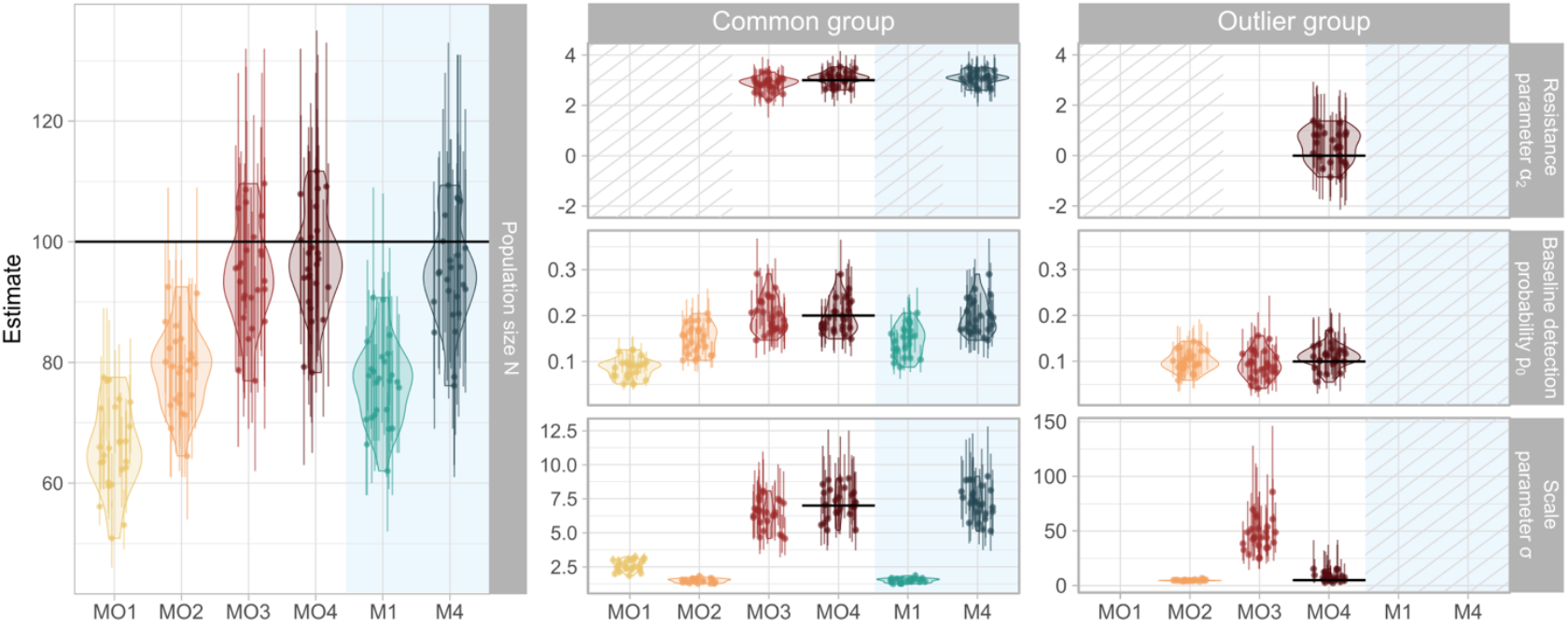
Model parameter estimates across the six models compared (see Table 1) in the simulation study. The mean estimate of each simulation is represented by a point with errors bar displaying the 95% credible intervals. Violin plots illustrate the distribution of the mean estimated parameters for each model across the 25 simulations. Horizontal lines indicate the true parameter values from model MO4, which was used to generate the simulated datasets. The first column shows population size estimates for the entire population. The second and third columns display parameter estimates for the common group and the outlier group, respectively. Within these columns, rows correspond to specific model parameters: the resistance parameter *α*_2_ (top row), baseline detection probability *p*_0_ (middle row), and scale parameter *σ* (bottom row). Background colors indicate data inclusion, with a white background representing models using all the dataset and a blue background indicating models with outliers removed. Striped areas denote parameters that were not estimated in the model. Empty spaces indicate cases where a single parameter was estimated for the entire population, with the reference parameter displayed in the common group.

#### 3.1.2 Connectivity estimates

Three models (MO3, MO4 and M4) allowed to look into connectivity estimates as they simultaneaously estimated individual and spatial heterogeneity in space use. Ignoring individual heterogeneity in the resistance parameter led to a resistance parameter close to the common group estimate that encompassed most of the population, with mean estimated *α*_2_ between 2.2 and 3.3 (MO3) compared to between 2.6 and 3.5 for the common group and between −0.9 and 1.4 for the outlier group (MO4) (Figure 3). By contrast, removing outlier indiduals led to results comparable to when individual and spatial heteorgeneity in space use was modelled, with mean estimated *α*_2_ between 2.6 and 3.5 (M4). To illustrate how these differences translate into the connectivity maps, we compared relative potential connectivity across these models (Figure 4), accounting for space use extent (*σ*) and its variability across the landscape (*α*_2_). Using simulated dataset 1 as an example, the three model formulations produced similar overall patterns. However, ignoring individual variation in the response to spatial heterogeneity in the landscape led to an overestimation of connectivity, especially visible in areas far away from the barrier (MO3 relative bias between −15% and 85.3%). Conversely, removing outliers led to an underestimation of connectivity close the barrier (M4 relative bias between - 86.7% and 26.1%).

**Figure 4.**
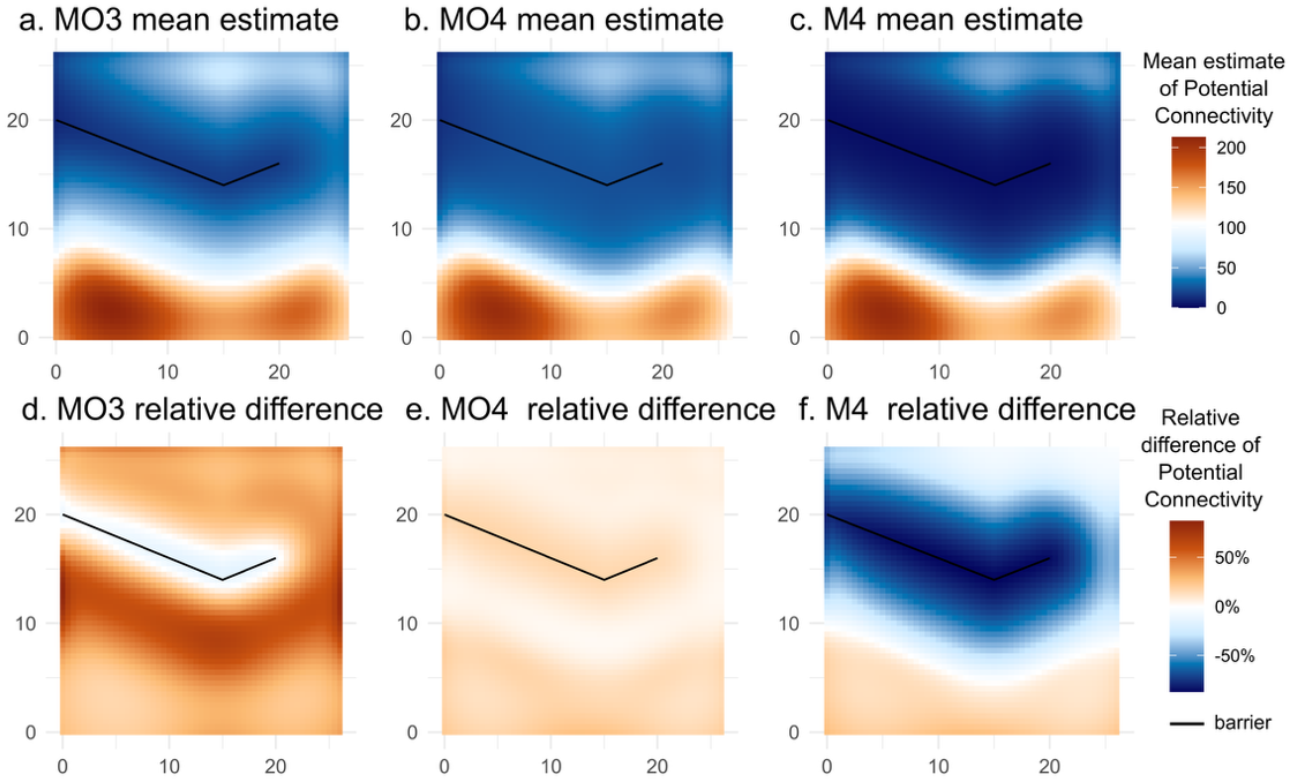
Illustration, for simulated dataset 1, of mean estimate (top row: a, b, c) and relative difference (bottom row: d, e, f) maps of potential connectivity. The left panels (a, d) show results for model MO3, which estimated a single resistance parameter *α*_2_for the entire population. The central panels (b, e) show results for model MO4, which estimated a separate resistance parameter *α*_2_for each group. The left panels (c, f) show results for model M4, where outlier individuals were removed from the analyses.

### 3.2 Case study

The models MFO1, MFO2, MFO4, MF1 and MF4 did converge with 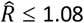 and effective sample size > 200. On the contrary, the mixing for the resistance parameter in the model MFO3 was poor with an effective sample size of only 93 and 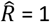.

#### 3.2.1 Population size estimates

In our case study, modelling individual heterogeneity in space use only (i.e. accounting that males are made of two groups, model MFO2) led to an increase of the estimated number of males (from 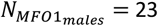 [20; 28] to 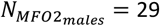 [21; 42] and 2 outliers) and a decrease of the estimated number of females (from 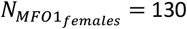 [85; 173] to 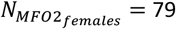 [50; 122]), compared to the null model (MFO1) (Figure 5). However, including spatial heterogeneity in space use did not affect population size estimates whether only one resistance parameter was estimated (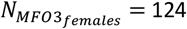 [80; 170], 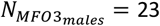 [20; 29] and 2 outliers) or one resistance parameter was estimated for each group (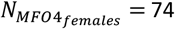 [47; 119], 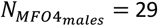 [21; 42] and 2 outliers).

**Figure 5.**
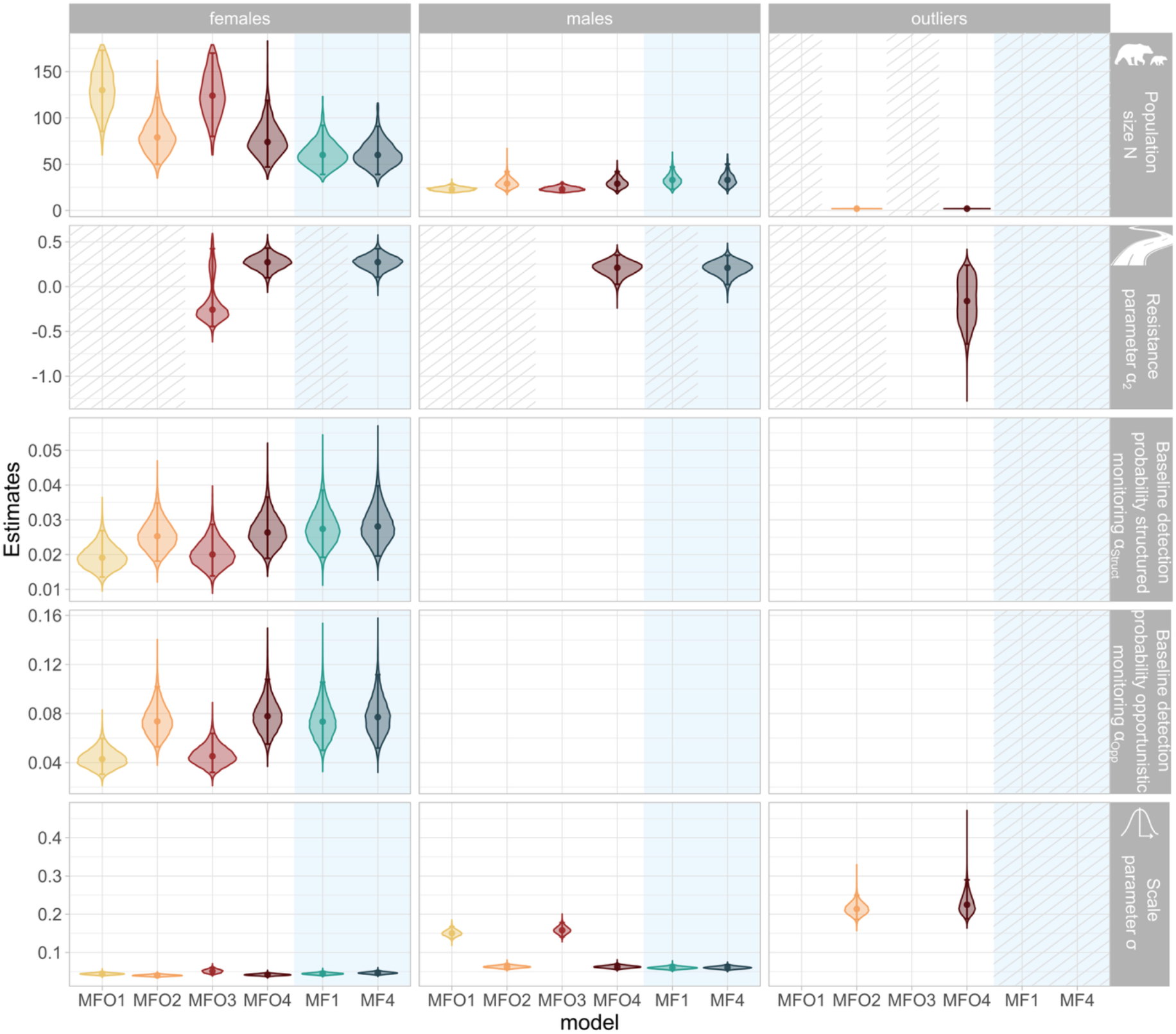
Violin plot of estimated population size N (top row), resistance parameter *α*_2_ (second row), intercept of baseline detection probability for the structured monitoring *α*_*Struct*_ (third row), intercept of baseline detection probability for the opportunistic monitoring *α*_0*pp*_ (fourth row), and scale parameter *σ* (bottom row) for the Pyrenean brown bear population. The median of each distribution is represented by a point with errors bar displaying 95% credible intervals. The panels in column display the estimated parameter”s posterior distribution for the females (left column), males (middle column) and outliers” (right column) Pyrenean brown bear population in 2019 in function of the six models compared. Background colours indicate data inclusion, with a white background representing models using all the dataset and a blue background indicating models with outliers removed. Striped areas denote parameters that were not estimated in the model. Empty spaces indicate cases where a single parameter was estimated for the entire population, with the reference parameter displayed in the female panel, except for the estimated distribution of the scale parameter in model MFO1 and MFO3 that is common to the males and the outliers and this parameter is therefore display in the male panel.

Removing the outliers from the dataset led to similar estimated population size than modelling individual heterogeneity (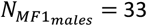 [23; 47], 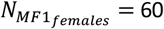 [39; 92]). Similarly, modelling spatial heterogeneity in space use when the outliers were removed led to comparable population size estimates (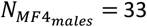 [23; 50], 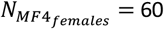 [39; 91]) (Figure 5).

The two models where outliers were removed had comparable estimates of population size (*N*_*MF*1_ = 93 [67; 131] to *N*_*MF*4_ = 94 [67; 133] (Figure 5)).

#### 3.2.2 Connectivity estimates

Integrating outlier individuals without modelling individual heterogeneity (model MFO1) led to a 2.7 times increase in the median estimated *σ* of males (*σ*_*MF*01_ = 0.15 [0.14; 0.17]), compared to when individual heterogeneity in space use was modelled (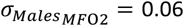 [0.06; 0.07]) or when removing outliers (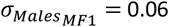 [0.05; 0.07]) (Figure 5).

Overall, the estimated resistance parameters was close to 0, whether we kept all the individuals in the dataset (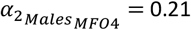 [0.03; 0.35], 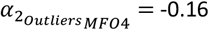 [-0.64; 0.24] and 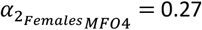 [0.01; 0.43]) or we removed the outliers (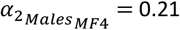 [0.02; 0.35] and 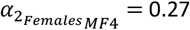 [0.11; 0.42] (Figure 5)). However, ignoring individual heterogeneity in the resistance parameter when keeping all the individuals in the dataset led to an estimated resistance parameter that encompassed 0 and increased parameter uncertainty (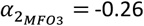 [-0.45; 0.42]) (Figure 5).

The map of density weighted connectivity predicted from model MFO4 showed that females” movements were more restricted than males” in high road density areas (Figure 6ac). However, the uncertainty of 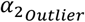 was propagated to the potential connectivity map, which did not allow us to draw any conclusion about their space use extent (Figure 6ef).

**Figure 6.**
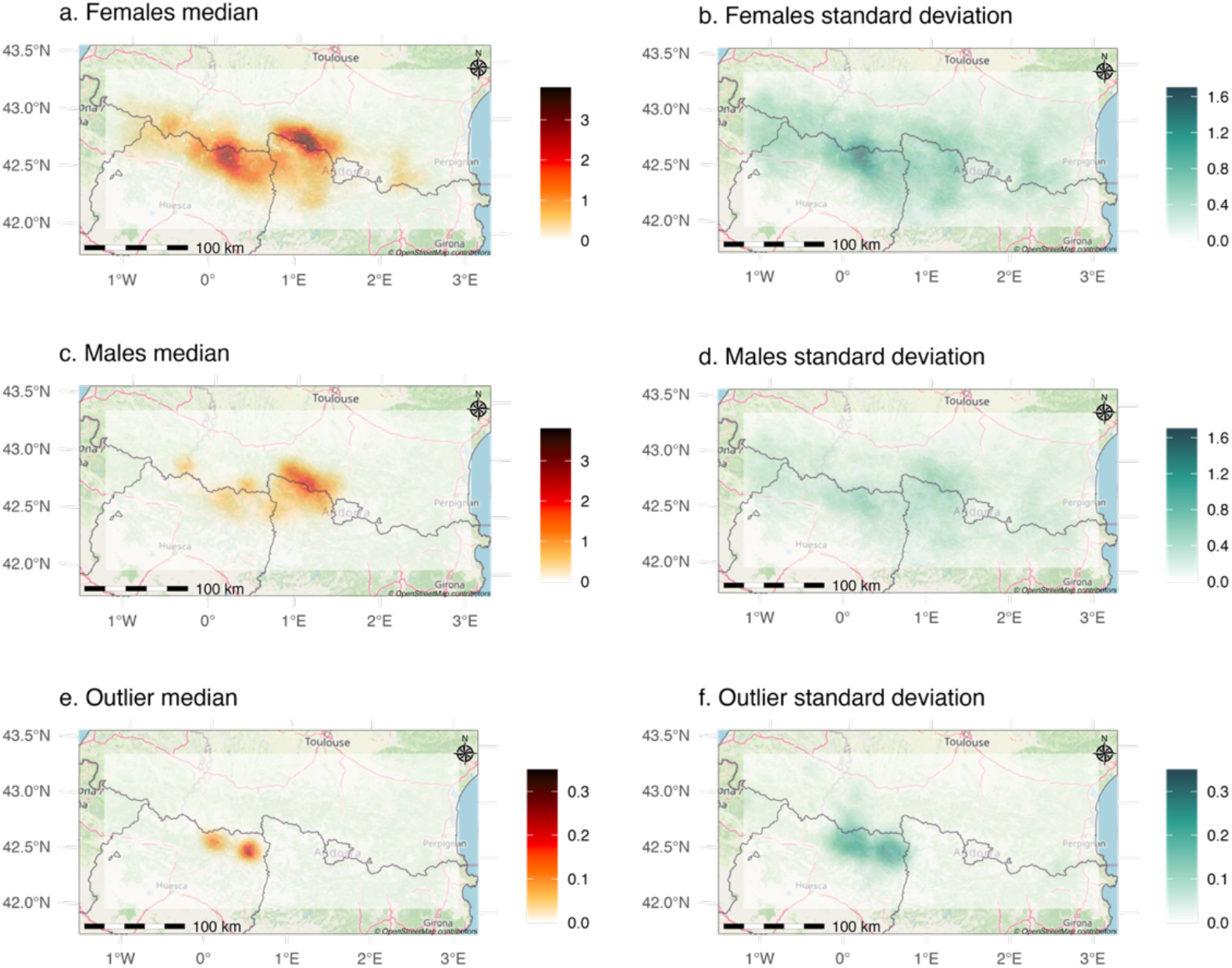
Maps of median density weighted connectivity derived from estimates of model MFO4, for females (a), males (b) and the two outlier individuals (e) with respective standard deviation (b, d, f) for the Pyrenean brown bear population in 2019.

## 4. Discussion

### 4.1. Population size estimates

As predicted, the simulation study revealed a negative bias in population size estimates when individual and spatial heterogeneities in space use were ignored (Pledger, 2000; Royle et al., 2013). This bias was strongest when both types of heterogeneity remained unmodelled.

Although modelling individual heterogeneity is challenging, especially when part of the population remains undetected (Marrotte et al., 2022; Pledger, 2000), we demonstrated how latent state formulations in SCR models can distinguish and estimate individual and spatial heterogeneity in space use. Conversely, when the primary objective is to estimate population size, removing outlier individuals with larger space use extent provides a straightforward way to reduce individual heterogeneity in space use and avoid underestimating population size (Kendall et al., 2019; Schmidt et al., 2022). However, this strategy is only effective if individual heterogeneity in space use is driven by a small number of individuals. Moreover, removing outlier individuals does not address spatial heterogeneity in space use that may remain in the population and would still require explicit modelling.

In our case study, understanding the combined effect of individual and spatial heterogeneity in space use on population size estimates was not straightforward. Applying the models to the Pyrenean brown bear population showed that individual heterogeneity could be accounted for, even with the low number of outliers (n=2). As expected, the estimated number of males was larger after accounting for individual heterogeneity (from 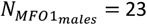 [20; 28] to 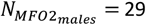 [21; 42] and 2 outliers). Surprisingly, the estimated number of females was smaller after accounting for individual heterogeneity (from 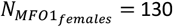 [85; 173] to 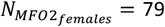 [50; 122]). In this study, we have not been able to unambiguously identify the reason of this observed pattern. When modelling individual heterogeneity, we noticed an increase in the intercept of the opportunistic baseline detection probability (from 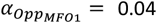 [0.03; 0.06] to 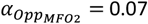 [0.05; 0.10]), while intercept of the structured baseline detection probability remained unchanged (from 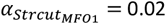 [0.01; 0.03] to 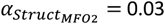 [0.02; 0.03]), linked to the presence of outliers in the population.

The estimated number of females was also larger than the 52 individuals detected in 2019 (with 22 males, 23 females and 7 individuals whose sex remains unknown)(Vanpé et al., 2022). A hypothesis to explain this apparent overestimation is that females have a smaller space use extent compared to males (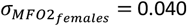 [0.036; 0.044] and 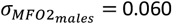 [0.057; 0.070]). From the model point of view, the small space use of female makes possible to miss them between traps, especially in areas that are covered solely by opportunistic traps. Additionally, in the south west part of the study area (Ordesa and Monte Perdido National Park), the monitoring intensity was low and even the habitat was perceived favourable (i.e. low human density and high ruggedness). This led to the prediction of high density of females in that area, which is unlikely given the low number of observed individuals in that area by the additional observations not included in this study. Unbiased estimation of population size will require working towards alternative model formulation and study design approaches to deal with those undesirable issues.

Furthermore, including spatial heterogeneity in space use in the model led to similar population size estimates compared to when only individual heterogeneity in space use was included (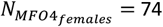 [47; 119], 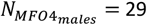 [21; 42] and 2 outliers). The estimated resistance parameter was close to zero for each group (respectively 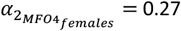 [0.10; 0.43], 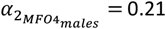 [0.03; 0.35] and 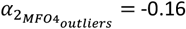 [-0.64; 0.24]) meaning that accounting for spatial heterogeneity in space use was not necessary in our case if the objective is to estimate population size (Sutherland et al., 2015).

Removing the outliers led to similar results that modelling individual heterogeneity (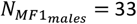 [23; 47], 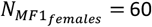 [39; 92]) demonstrating that this practice is a cost effective solution when the aim was to estimate population size (Kendall et al., 2019; Schmidt et al., 2022).

### 4.2 Connectivity estimates

The resistance parameter reflects whether space use extent increases (*α*_2_ < 0) or decreases (*α*_2_ > 0) as a response to a landscape covariate (Royle et al., 2013; Sutherland et al., 2015). In our simulations, the common group had smaller space use extent close to the barrier, while the outlier group had larger space use extent and was not affected by landscape structure. Under these conditions, the model that ignored the individual heterogeneity in the resistance parameter (model MFO3) estimated a resistance parameter skewed toward the common group value, suggesting that all individuals in the population reduced space use near the barrier. A comparable conclusion would be drawn when removing outlier individuals from the data set with a single estimated resistance parameter.

Potential connectivity maps enable to map population space use by assuming an activity centre in each cell of the spatial domain and summing the space use of each individual (Morin et al., 2017). Comparing maps from models MFO3, MFO4 and MF4 showed consistent reductions in connectivity near barriers, but values were sensitive to model specification (Figure 4). Ignoring individual heterogeneity in the resistance parameter (model MFO3) led to overestimating potential connectivity further away from the barrier, whereas removing outliers (model MF4) led to underestimating potential connectivity at the barrier. These results highlight the need for caution when comparing potential connectivity maps.

In the case study, the estimated resistance parameter demonstrated that both males and females exhibited smaller space use extents in areas with higher road density, and their responses were similar across sexes (Figure 5).

Density Weighted Connectivity (DWC) maps provide an integrated representation of population space use by combining realized densities, individual heterogeneity in space use extent (scale parameter *σ*), and heterogeneous responses to landscape features (resistance parameter *α*_2_). These maps allow delineation of areas with high space use as well as those where accessibility is limited by the road network. In addition, the uncertainty maps associated with DWC showed that the outlier group map uncertainty is too high to support reliable conclusions, while the map for females identifies in Ordesa and Monte Perdido National Park, elevated uncertainty that suggests the need for further investigation in this area to refine space use assessment (Figure 6).

DWC maps are based on realized densities which are known to be sensitive to the location of traps in the landscape (Durbach et al., 2024), contrary to predicted maps that are based on the estimated relationship between density and landscape variables. Given the difficulty to describe spatial variation in the Pyrenean brown bear density based on spatial covariates (Sanz-Pérez et al., 2025), and considering the intensive monitoring of the population, it is likely that the realized densities more accurately reflect the population distribution than predicted maps.

Overall, the effects of individual and spatial heterogeneity in space use on connectivity estimates are much more challenging to interpret than that on population size: they are sensitive to model formulation, trap location, and there is no clear parameter comparison to validate the outputs. Thus, interpreting SCR-based connectivity requires caution, even though jointly modelling spatial and individual heterogeneity remains conceptually appealing.

### 4.3 Challenges and perspectives

Our findings highlight the importance of tailoring the structure of SCR model to the research objectives. When the objective is to estimate population size, removing outlier individuals is an efficient solution that avoids unnecessary complexity. When the goal is to assess connectivity, however, disentangling individual and spatial heterogeneity is essential but more challenging.

Identifying outliers is a first challenge and itself subjective. In our case, we examined the distribution of the capture frequency and the maximum distance moved (MDM) as indicators, both of which can reveal unusual detectability patterns. While further research is necessary to identify outliers, these summary statistics are a good first approach to detect obvious outliers (Figure 2, Table 2). In addition, comparing population size estimates with the presence and absence of outliers may be a good test to decide whether accounting for individual heterogeneity is necessary. Large differences across models indicate that results are sensitive to heterogeneity assumptions, underscoring the importance of testing alternative formulations.

**Table 2.** Number of females, males and outliers detected and number of detections across traps and sampling occasions each year (2017, 2018, 2019) by the structured and the opportunistic monitoring in the Pyrenean brown bear population.

|  | females | males | outliers |
| --- | --- | --- | --- |
| **Number of individuals detected** |  |  |  |
| 2017 | 17 | 14 | 2 |
| 2018 | 14 | 17 | 2 |
| 2019 | 16 | 17 | 2 |
| **Number of detections** |  |  |  |
| 2017 | 47 | 53 | 30 |
| 2018 | 32 | 57 | 32 |
| 2019 | 58 | 89 | 22 |

**Table 3.** Models formulas used for the simulation study and applied on each simulated datasets. It present the model name, the dataset, the scale parameter (σ), the resistance parameter (α_2_) when applicable. The empty cells means that the resistance parameter was not included in the model and therefore we used the Euclidean distance instead of the least-cost path distance.

| Model name | dataset | $\sigma$ | $\alpha_2$ |
| --- | --- | --- | --- |
| MFO1 | all | ~sex |  |
| MFO2 | all | ~group |  |
| MFO3 | all | ~sex | ~1 |
| MFO4 | all | ~group | ~group |
| MF1 | outlier removed | ~sex |  |
| MF4 | outlier removed | ~sex | ~sex |

Accounting for individual heterogeneity also depends on data availability (Schmidt et al., 2022). In this study, we assumed known group membership for detected individuals and used latent state formulations to assign undetected individuals. When group membership is unknown, finite mixture models can be applied (Marrotte et al., 2022; Pledger, 2000), though interpretation of latent groups requires caution (Pledger and Phillpot, 2008). We provide the code for the finite mixture SCR model with ecological distance in *nimble* when groups are unknown (Appendix S1).

## 5. Conclusion

Our study underscores that SCR models must be tailored to research goals. When the objective is to estimate population size, removing outliers can provide efficient and unbiased estimates without increasing model complexity. When the objective is to assess connectivity, however, explicitly modelling both spatial and individual heterogeneity is necessary, despite the challenges. Ultimately, model complexity should be matched to both data availability and the ecological questions at hand to ensure robust and meaningful inference for management and conservation.

## Acknowledgements

We gratefully acknowledge all brown bear field co-workers from the Brown Bear Network (ROB) and from Spanish and Andorran teams. We thank Perry de Valpine for his help on the implementation of the model in Nimble. The simulations were performed on the Core Cluster of the Institut Français de Bioinformatique (IFB) (ANR-11-INBS-0013). We thank the two referee, for their helpful and constructive comments.

## Funding

The work was partially supported by the French National Research Agency grant ANR-16-CE02-0007. Financial and logistical supports for this study were provided by the ONCFS / OFB in France. Genetic analyses were performed by the ANTAGENE Company in France and the Servei Veterinari de Genètica Molecular, Universitat Autònoma de Barcelona in Spain. This research was partly funded by Biodiversa+, the European Biodiversity Partnership, in the context of the Big_Picture project under the 2022-2023 BiodivMon joint call. It is co-funded by the European Commission (GA No. 101052342) and the French Agence Nationale de la Recherche.

## Conflict of interest disclosure

The authors declare no conflict of interest.

## Data, scripts and supplementary information availability

Appendices, scripts and simulations data are available on the online Zenodo repository (https://doi.org/10.5281/zenodo.17259654). Pyrenean brown bear detection data as well as trap locations are sensitive data and are therefore not provided in this repository.

## Notes

https://doi.org/10.5281/zenodo.17259654

### Competing Interest Statement

The authors have declared no competing interest.

### Summary of Updates

This version has been recommended by PCI Ecology

