## Appendix S1 for "Unravelling the effects of heterogeneity in space use on estimates of connectivity and population size: Insights from spatial capture-recapture modelling"

### Appendix S1: Unravelling the effects of heterogeneity in space use on estimates of connectivity and population size: Insights from spatial capture-recapture modelling

```
source("SimulationsV2/functions/AccessCachedNim.R")
```

M. Kervellec, O. Gimenez, C. Vanpé, P.-Y. Quenette, J. Sentilles, S. Palazón, I. Afonso Jordana, R. Jato, M. M. Elósegui Irurtia, and C. Milleret

In the following appendix we provide the script to simulate spatial detection data for two groups and the nimble model. First we assume that groups are known as presented in the manuscript and then we provide an example to run the model when groups are unknown (mixture model)

### 1 Simulate data

#### 1.1 Population parameters

```
# Group a
N.a <- 97 # population size
p0.a <- 0.2 # baseline detection probability
sigma.a <- 7 # scale parameter
alpha1.a <- 1/(2*sigma.a*sigma.a)
alpha2.a <- 3 # resistance parameter

# Group b = outliers
N.b <- 3
DeltaP0 <- -0.1
p0.b <- p0.a + DeltaP0
DeltaSigma <- -2
sigma.b <- sigma.a + DeltaSigma
alpha1.b <- 1/(2*sigma.b*sigma.b)
DeltaAlpha <- -3
alpha2.b <- alpha2.a + DeltaAlpha

N <- N.a + N.b
```

#### 1.2 Study Area and resistance covariate

```
# 1. Constant definition -------------------------------------------------------
K <- 7 # number of occasions
Array <- 10 # number of traps in one line: size of the trap array
```

```r
Resolution <- 0.5 # resolution of the resistance surface
trap_spacing <- 2 # 2*sigma.a

# 2. State space ----------------------------------------------------------------
# Set a squared study area according to recommendation
buff <- 4*(trap_spacing/2) # size of the buffer -> 4*sigma
L <- 2*buff*(1/Resolution) + (Array-1)*trap_spacing*(1/Resolution) # total number of cell in a row

# 3. Trap locations -------------------------------------------------------------
traplocs <- expand.grid(X = seq(buff,
                                (L*Resolution)-buff,
                                trap_spacing),
                        Y = seq(buff,
                                (L*Resolution)-buff,
                                trap_spacing)) %>% # trap spacing = 2*sigma
  as.matrix()
ntraps <- nrow(traplocs)

# 4. Activity center location: Homogeneous point process -----------------------
spatialdomain <- expand.grid(X = seq(0, (L+1)*Resolution,1),
                             Y = seq(0, (L+1)*Resolution,1)) %>%
  as.matrix()
# sample across the locations in the spatial domain
S.a <-tibble(X = sample(spatialdomain[,1], size = N.a, replace = TRUE),
             Y =  sample(spatialdomain[,2], size = N.a, replace = TRUE)) %>%
  as.matrix()


S.b <-tibble(X = sample(spatialdomain[,1], size = N.b, replace = TRUE),
             Y =  sample(spatialdomain[,2], size = N.b, replace = TRUE)) %>%
  as.matrix()

S <- rbind(S.a,S.b) %>%
  as.data.frame() %>%
  mutate(group = c(rep("a",nrow(S.a)),rep("b",nrow(S.b))),
         ind = row_number())

# 5. Define resistant surface covariate ----------------------------------------
covariate <- expand_grid(seq(0,L*Resolution,Resolution),
                         seq(0,L*Resolution,Resolution)) %>%
  mutate(z  = runif((L+1)^2, -1, 1)) %>%
 rasterFromXYZ()

# set.seed(2025)
# covariate <- predict(gstat(formula = z ~ 1,
#                            locations = ~ x + y,
#                            dummy=T, beta=1,
#                            model=vgm(psill=1,model="Exp",range=20),
#                            nmax=20), # define the gstat object (spatial model)
#                      newdata = grid %>%
#                        raster::as.data.frame(xy = TRUE), # make one simulations based on the stat obj
#                      nsim = 1) %>%
#   mutate(sim1 =((sim1 - min(sim1))/(max(sim1)-min(sim1)))) %>% # scale between 0 & 1
```

```r
#   raster::rasterFromXYZ()

x_coords <- c(0,15,20)
y_coords <- c(20, 14, 16) # define the line points

points <- sp::SpatialPoints(cbind(x_coords, y_coords)) %>% # from spatial points
  sf::st_as_sf() %>%
  sf::st_combine()%>%
  sf::st_cast("LINESTRING") # convert to line

covariate[!is.na(covariate[])] <- sf::st_distance(SpatialPoints(coordinates(covariate)[!is.na(covariate
                                        points, byid = T) # from each resistance raster cell, we comput
covariate <- 1-((covariate - min(raster::values(covariate)))/(max(raster::values(covariate))-min(raster

# 6. look at what we have created so far ---------------------------------------
ss.plot <- ggplot() +
  geom_raster(covariate %>% raster::as.data.frame(xy=TRUE),
              mapping = aes(x = x, y = y, fill = z)) +
  scale_fill_distiller(palette = "Spectral", direction = -1) +
  geom_point(S, mapping = aes(x=X, y = Y, color = group))+
  geom_point(traplocs %>% as.data.frame(), mapping = aes(x=X, y = Y), pch = 3) +
  theme_light()

ss.plot +
  ggtitle("Distribution of traps, activity centers on the resistance covariate")
```

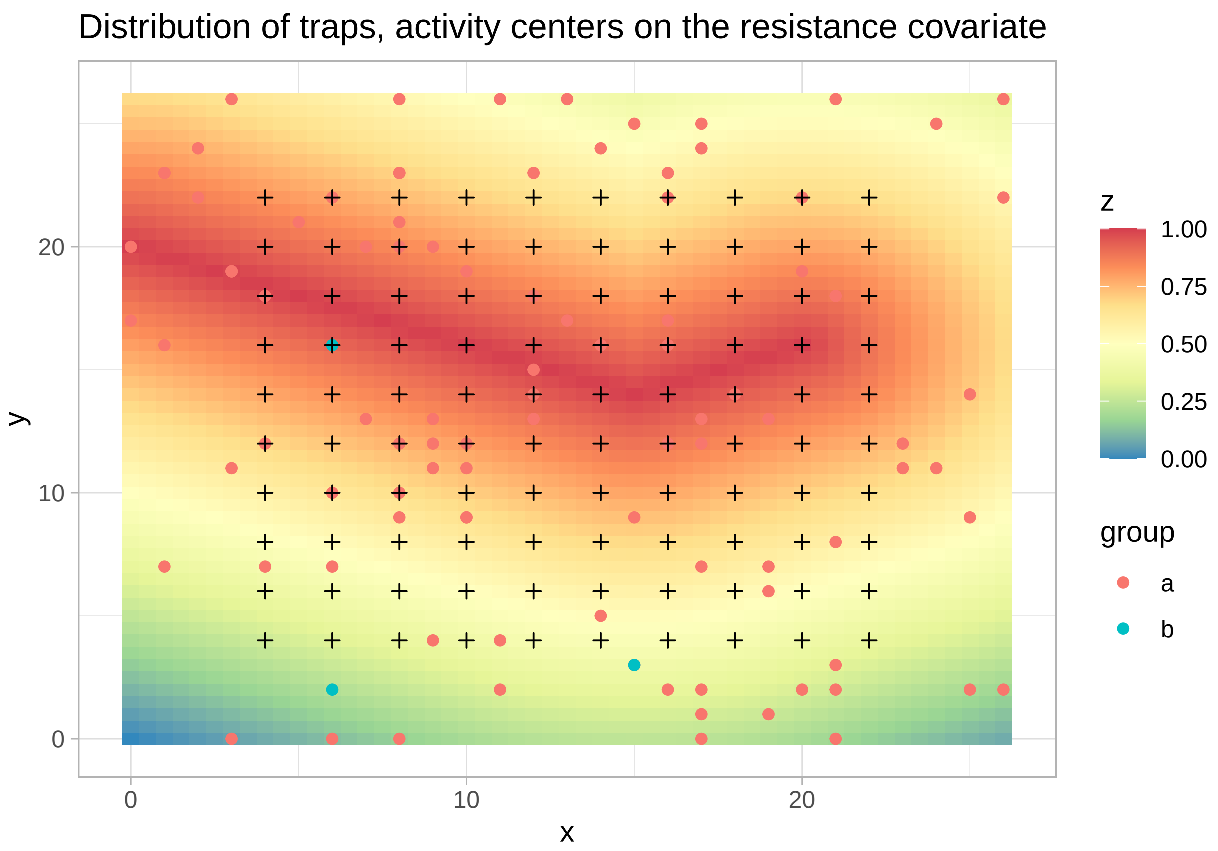

In this figure th underlying raster represents the length of roads per pixel. The black crosses represent traps and the points the activity centers simulated according to an homogeneous point process for group "a" in red and group "b" in blue.

##### 1.3 Simulate Detections

Define resistance scenarios for landscape A Find a sigma for a given alpha2

```
source("SimulationsV2/functions/mean_homerange_size.R")
## extract coefficients  dont load this function it already exist in secr
coeff <- function(alpha2, data){
  data.a2 <- data %>%
    filter(a2 == alpha2) %>%
    mutate(sigma2 = sigma^2)
  lm <- lm(data.a2$HR ~ data.a2$sigma + data.a2$sigma2)
  return(c(coefficients(lm),alpha2))}

# solve polynomial, i.e. find the sigma given alpha and home range size
solve.poly <-function(alpha2, coef,HRsize){
  row <- which(coef[,4] == alpha2) # retrieve the coefficients according to a given alpha2
  Re(polyroot(c(coef[row,1]-HRsize, coef[row,2],coef[row,3])))[which(Re(polyroot(c(coef[row,1]-HRsize, c
}

# Define set of alpha2 and sigma by hand
SC <- data.frame(a2 = c(rep(0,12),
                        rep(1,6),
                        rep(2,12),
                        rep(3,10),
                        rep(4,10),
                        rep(5,13)),
                 sig = c(seq(0.1, 1.2, by=0.1),
                         seq(0.1, 3, by=0.5),
                         seq(0.1,6, by=0.5),
                         seq(0.1, 10, by=1),
                         seq(0.1, 20, by=2),
                         seq(0.1, 25, by=2)))

data <- data.frame(a2 = NA,
                   HR  = NA,
                   sigma = NA) # initialize output matrix

# Compute mean HR
for (i in 1:nrow(SC)){
  #print(i)
  out <- mean_homerange_size(sigma = SC[i,2],
                             a2 = SC[i,1],
                             covariate = covariate,
                             x.box = c(4,22), # where the HR are computed
                             y.box = c(4,22))
  data[i,] <- out
}

# Fit an exponential model to each of the curve
```

```
Coefficients.poly <- coeff(0, data = data) %>%
  rbind(coeff(1, data = data)) %>%
  rbind(coeff(2, data = data)) %>%
  rbind(coeff(3, data = data)) %>%
  rbind(coeff(4, data = data)) %>%
  rbind(coeff(5, data = data))

# For each alpha for a set Home Range size find the corresponding sigma (choose HR such as sigma = 1 fo
HomeRangeSize <- 105
Coefficients.model <- data.frame(sigma = c(solve.poly(0, coef = Coefficients.poly, HRsize = HomeRangeSi
                                  solve.poly(1, coef = Coefficients.poly, HRsize = HomeRangeSize),
                                  solve.poly(2, coef = Coefficients.poly, HRsize = HomeRangeSize),
                                  solve.poly(3, coef = Coefficients.poly, HRsize = HomeRangeSize),
                                  solve.poly(4, coef = Coefficients.poly, HRsize = HomeRangeSize),
                                  solve.poly(5, coef = Coefficients.poly, HRsize = HomeRangeSize)),
                        a2 = c(0:5))

# Have a look to the results
row <- 5
sig.sets <- seq(0,1.2,0.5)
Check <- data.frame(sig = sig.sets,
                    HR = Coefficients.poly[row,1] + Coefficients.poly[row,2]*sig.sets + Coefficients.pol
HRsizeplot <- ggplot() +
  geom_point(data, mapping = aes(x = sigma, y = HR, col = as.factor(a2))) +
  geom_line(data, mapping = aes(x = sigma, y = HR, col = as.factor(a2))) +
  geom_point(Coefficients.model, mapping = aes(x = sigma, y  = HomeRangeSize, col = as.factor(a2)), pch
  geom_hline(yintercept = HomeRangeSize) +
  theme_light()
  #geom_line(Check, mapping = aes(x = sig, y = HR))
HRsizeplot
```

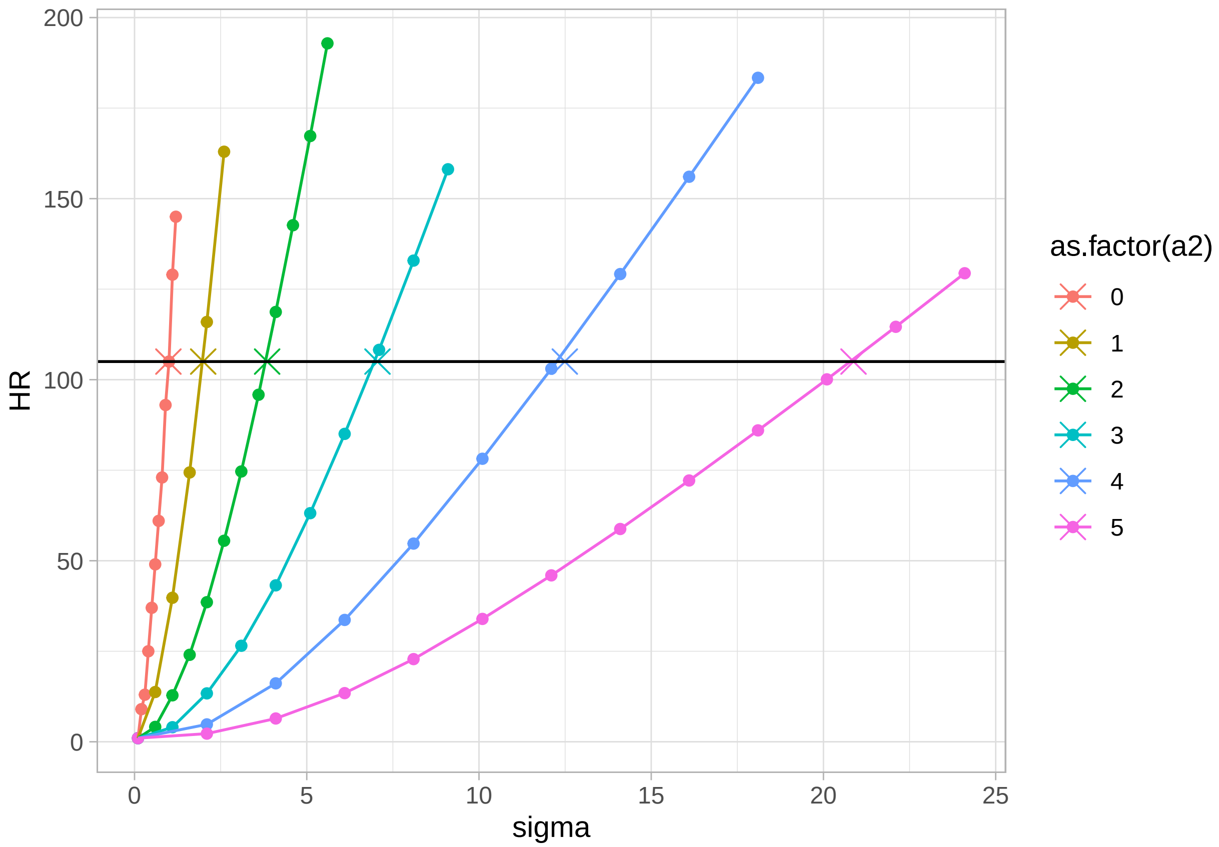


```
#ggplot2::ggsave(plot = HRsizeplot, dpi = 600,  width = 4.5, height = 4, filename = "plots/LandscapeAHR

# We define accordingly a sigma for each alpha2
Coefficients.model
```

```
##        sigma a2
## 1  0.9829354  0
## 2  1.9931158  1
## 3  3.8496701  2
## 4  7.0527836  3
## 5 12.4913487  4
## 6 20.8730084  5
```

```
alpha2 <- c(0, 1, 2, 3, 4, 5)
sigma <- c(1, 2, 4, 7, 12, 21)
```

```
# 1. Compute detection probabilities for each individual -----------------------
## group a
cost.a <- exp(alpha2.a * covariate) # compute cost of each cell

D.a <- gdistance::transition(cost.a,
                             transitionFunction=function(x) 1/mean(x),
                             directions=8) %>%
  gdistance::geoCorrection(type="c",multpl=F,scl=F) %>%
  gdistance::costDistance(S.a, traplocs) # compute LCP distance between AC and traps, same as e2dist(S,
```

```
probcap.a <- p0.a * exp(-alpha1.a * D.a * D.a) # compute detection probabilities

## group b
cost.b <- exp(alpha2.b * covariate)

D.b <- gdistance::transition(cost.b,
                             transitionFunction=function(x) 1/mean(x),
                             directions=8) %>%
  gdistance::geoCorrection(type="c",multpl=F,scl=F) %>%
  gdistance::costDistance(S.b, traplocs)

probcap.b <- p0.b * exp(-alpha1.b * D.b * D.b)

probcap <- rbind(probcap.a, probcap.b)

# 2. Simulate detection according to defined detection probabilities -----------
y <- array(0, dim=c(N, ntraps, K)) # empty 3D array (individuals by traps by sampling occasion)

# Simulate encounter histories
for(i in 1:N){                                # loop through each individual/activity center
  for(j in 1:ntraps){                         # loop through each trap
    y[i,j,1:K]<- rbinom(K, 1, probcap[i,j])  # y ~ binomial(p_ijk)
  }
}

Y <- apply(y, c(1,2), sum) # number of time an individual was captured at a trap in a session
#Y
caps.per.ind <- apply(y,1,sum) # shows # captures for each individual across all traps and all occasion
#caps.per.ind
N.det <- length(caps.per.ind[which(caps.per.ind>0)])
#N.det

# Keep detection history only for individuals detected
M <- N + 50 # augmented population size

eta_det <- c(rep(0, N.a),rep(1, N.b))[which(caps.per.ind>0)]
eta_data <- c(eta_det, rep(NA, M-N.det))

# Augmenting y
yy <- array(rep(0,M*ntraps*K), c(M,ntraps,K)) # augmented detection array
yy[1:N.det,,] <- y[which(caps.per.ind>0),,]

# 3. look at encounter probabilities of a few individual -----------------------
someGuys.a <- S.a[sample(1:N.a,5),]
D2.a <- gdistance::transition(cost.a,
                              transitionFunction=function(x) 1/mean(x),
                              directions=8) %>%
  gdistance::geoCorrection(type="c",multpl=F,scl=F) %>%
  gdistance::costDistance(someGuys.a, raster::coordinates(covariate))
probcap2.a <- p0.a * exp(-alpha1.a * D2.a * D2.a)

someGuys.b <- S.b[sample(1:N.b,3),]
D2.b <- gdistance::transition(cost.b,
```

```
                                      transitionFunction=function(x) 1/mean(x),
                                      directions=8) %>%
  gdistance::geoCorrection(type="c",multpl=F,scl=F) %>%
  gdistance::costDistance(someGuys.b, raster::coordinates(covariate))
probcap2.b <- p0.b * exp(-alpha1.b * D2.b * D2.b)

ggplot()+
  # group a
  geom_raster(data.frame(raster::coordinates(covariate),
                         detectionProb = apply(probcap2.a,2,sum)),
              mapping = aes(x=x, y = y, fill = detectionProb)) +
  scale_fill_distiller(palette = "Reds", direction = 1) +
  geom_point(someGuys.a %>% as.data.frame() , mapping = aes(x=X, y = Y)) +
  # traps
  geom_point(traplocs %>% as.data.frame(), mapping = aes(x=X, y = Y), pch = 3) +
  theme_light() +
  ggtitle("Detection probability of 5 individuals of group a ")
```

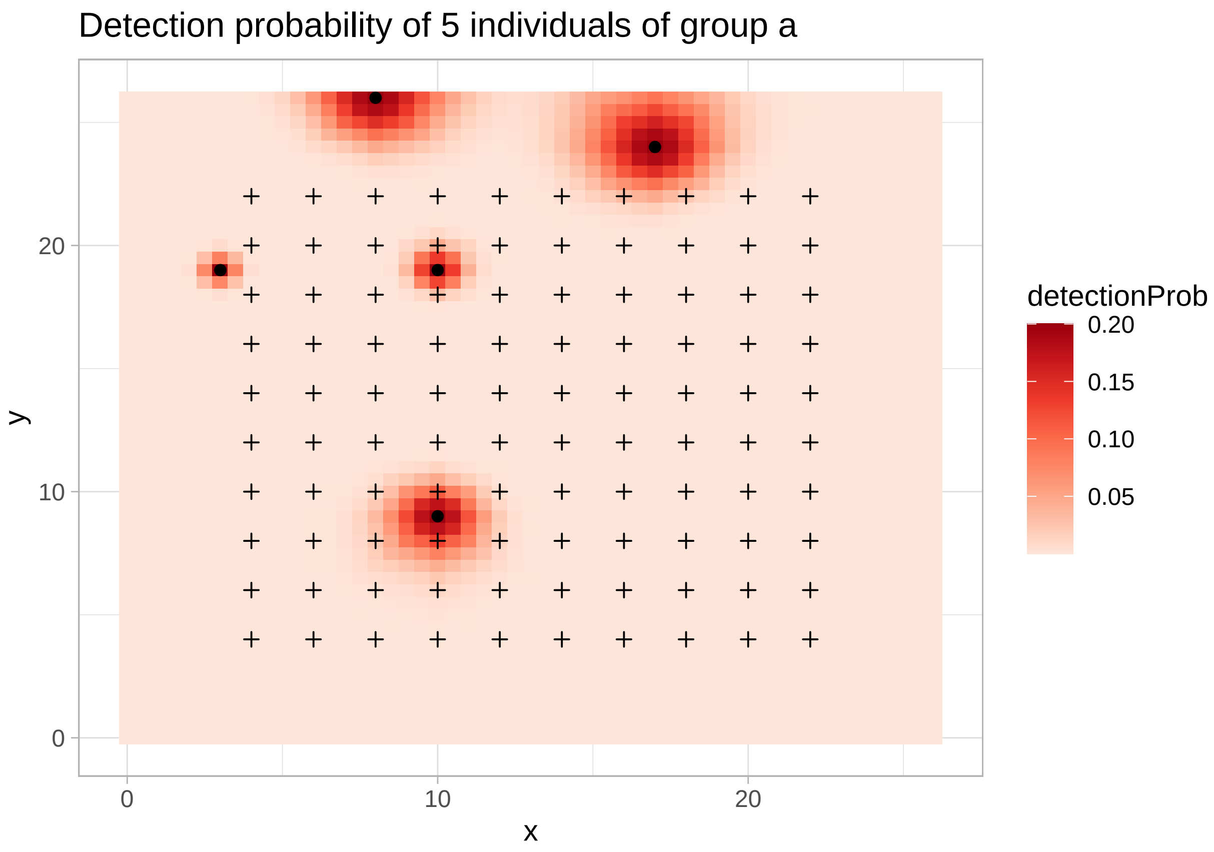


```
ggplot()+
  # group b
  geom_raster(data.frame(raster::coordinates(covariate),
                         detectionProb = apply(probcap2.b,2,sum)),
              mapping = aes(x = x, y = y, fill = detectionProb)) +
  scale_fill_distiller(palette = "Blues", direction = 1) +
  geom_point(someGuys.b %>% as.data.frame() , mapping = aes(x=X, y = Y)) +
```

```
  # traps
  geom_point(traplocs %>% as.data.frame(), mapping = aes(x=X, y = Y), pch = 3) +
  theme_light() +
  ggtitle("Detection probability of 3 individuals of group b")
```

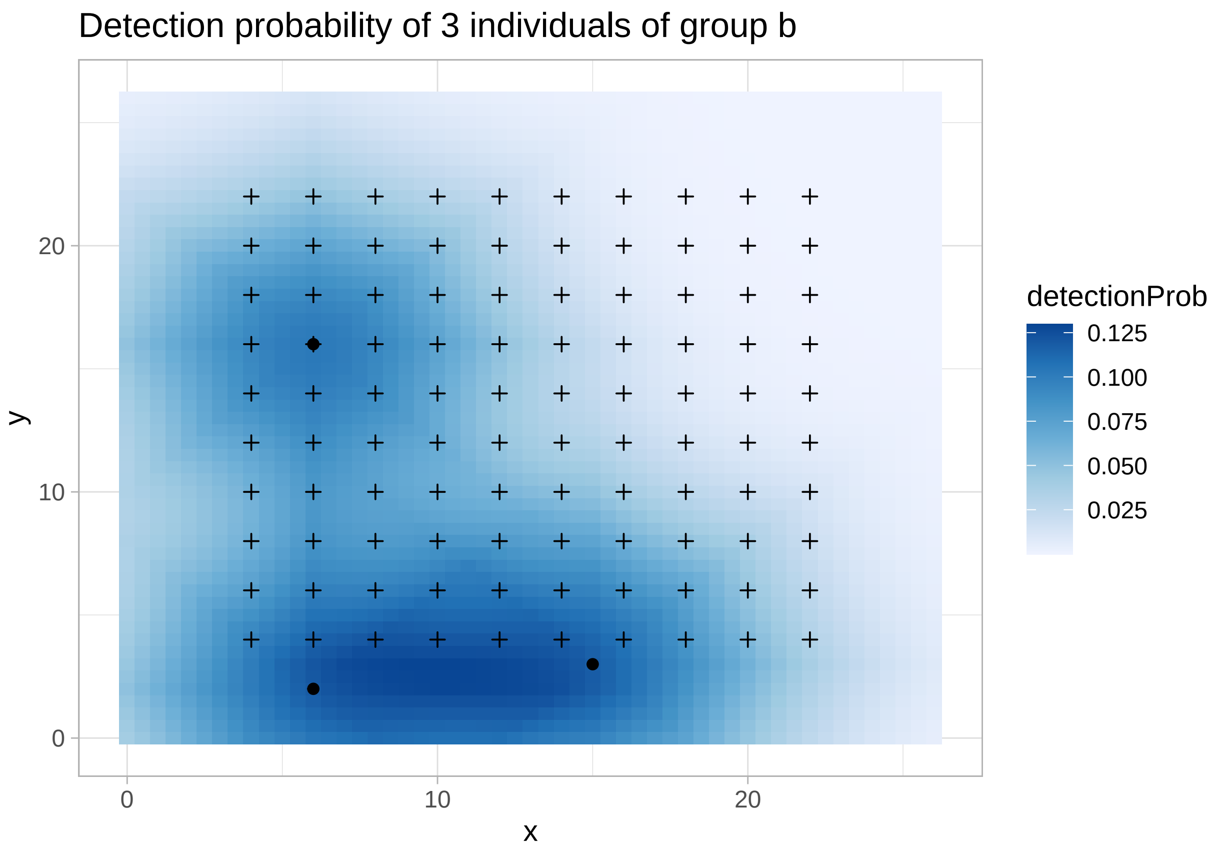


These two maps represents the cumulative detection probability across the state space for the five randomly chosen activity center respectively for group "a" and group "b"

```
# 4. look at detection histories -----------------------------------------------
tdf <- traplocs %>%
  as.data.frame() %>%
  mutate(trap = paste0("V",c(1:ntraps)) %>%
           as.factor() %>%
           as.numeric())

for (k in 1:K){
  Det <- y[,,k] %>% # matrix of detections
    as_tibble(rownames = "ind", colnames = "trap") %>%
    pivot_longer(cols = -ind, names_to = "trap", values_to = "occ") %>% # transform to a longer datafra
    mutate(trap = as.numeric(as.factor(trap))) %>%
    filter(occ > 0) %>% # We only keep detections
    mutate(occ = occ*k) %>% # identify the occasion of capture
    left_join(tdf) %>% # retrieve trap location
    left_join(S %>% # retrieve AC location and group of each ind
                as.data.frame() %>%
                mutate(ind = as.character(1:nrow(S))) %>%
```

```
                rename(sx=X,
                       sy=Y),
              by = "ind")
  # save for each occasion
  if(k==1){
    Dets <- Det}
  else{
    Dets <- rbind(Dets,Det)}
}
```

```
## Warning: The `x` argument of `as_tibble.matrix()` must have unique column names if
## `.name_repair` is omitted as of tibble 2.0.0.
## i Using compatibility `.name_repair`.
## This warning is displayed once every 8 hours.
## Call `lifecycle::last_lifecycle_warnings()` to see where this warning was
## generated.
```

```
## Joining with `by = join_by(trap)`
## Joining with `by = join_by(trap)`
## Joining with `by = join_by(trap)`
## Joining with `by = join_by(trap)`
## Joining with `by = join_by(trap)`
## Joining with `by = join_by(trap)`
## Joining with `by = join_by(trap)`
```

```
# Plot
ss.plot +
  geom_segment(data=Dets,
               aes(x=sx,y=sy,xend=X,yend=Y, color = group),lwd=0.3) +
  geom_point(S %>% as.data.frame() %>%  mutate(ind = as.character(1:nrow(S))) ,
             mapping = aes(x=X, y = Y, color = group)) +
  ggtitle("Links between detections and activity centers")
```

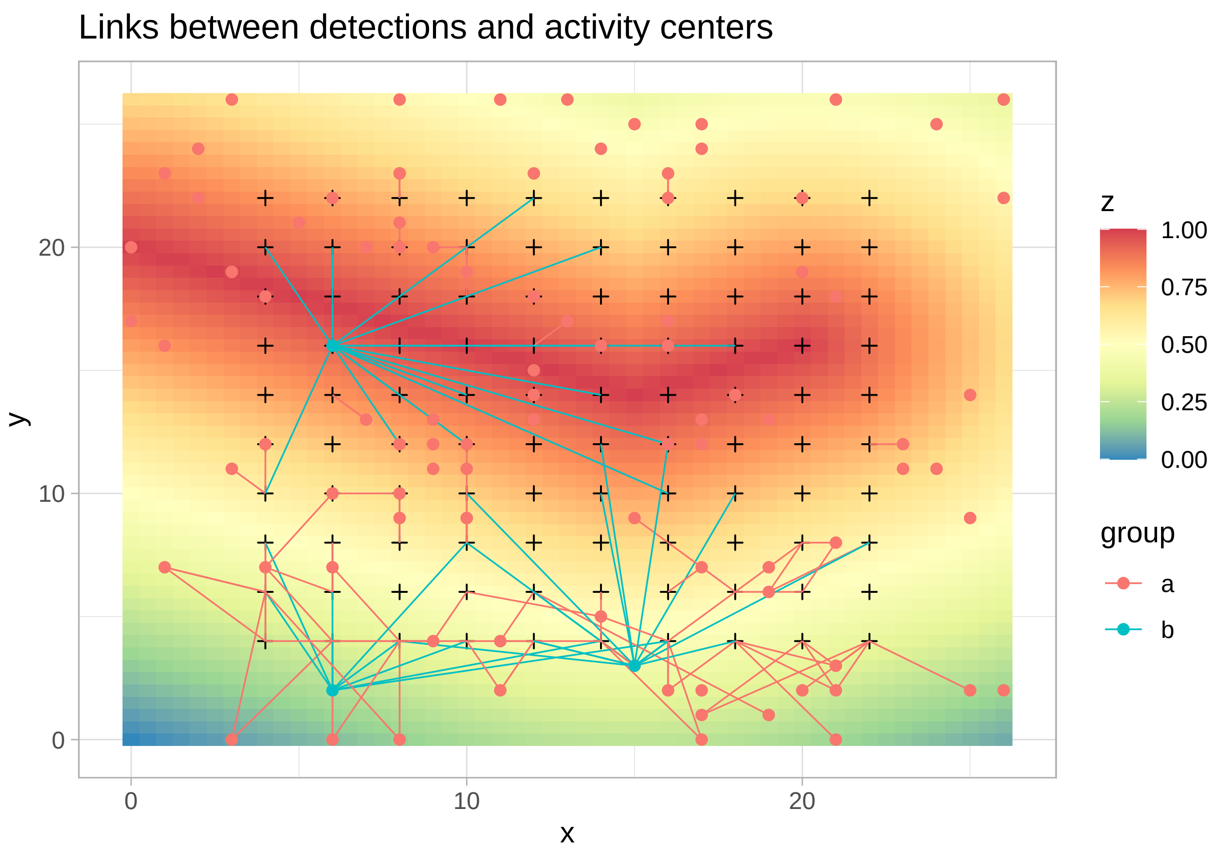


Then, this figure shows where each individual was detected by drawing lines between simulated activity centers and the traps where the individual was detected.

### 2. Fit model

#### 2.1 Model with 2 known groups

```
# Least cost path function
leastcostpath <- function(alpha){
  # alpha is a scalar
  # rcov is a RasterLayer
  # xy is a matrix of dims nsites x 2
  # obtain resistance surface
  cost <- exp(alpha * covariate)

  D <- gdistance::transition(cost, transitionFunction=function(x) 1/mean(x), directions=8) %>%
    gdistance::geoCorrection(type="c",multpl=F,scl=F) %>%
    gdistance::costDistance(spatialdomain, traplocs)

  D^2
}

# Convert the function into NIMBLE
```

```
LCP <- nimbleRcall(function(alpha = double(0)){},
                   Rfun = 'leastcostpath',
                   returnType = double(2))

A <- leastcostpath(-5) # test
```

Compute cached matrix

```
# make correspondance between cell corrdinates and rownumber in the distance array
habitatgrid <- matrix(1:nrow(spatialdomain), nrow = max(spatialdomain[,1])+1) # identify AC location

cells = nrow(spatialdomain) # number of cells in the spatial domain
x.max = y.max = L*Resolution # extend of the spatial domain

# Alpha array
step_alpha <- 0.1
alphaValues <- round(seq(-5, 5, by=step_alpha), digits=1)

cached <- array(NA, c(cells, ntraps, length(alphaValues)))

for(a in 1:length(alphaValues)){
  cached[1:cells,1:ntraps,a] <- LCP(alphaValues[a]) # compute distances between habitat cells and traps
}

roundValue <- 1/step_alpha
indexValue <- which(alphaValues<=0) %>% length()
```

Model Code

```
modelCode2groups <- nimbleCode({
  ## priors
  psi ~ dunif(0, 1)
  probB ~ dunif(0, 1)

  sigma[1] ~ dunif(0, 50)
  DeltaSigma ~ dunif(-50, 50)
  sigma[2] <- sigma[1] +  DeltaSigma
  three ~ dconstraint(sigma[2] > 0) # positive scale parameter

  p0[1] ~ dunif(0, 1)
  DeltaP0 ~ dunif(-1, 0)
  p0[2] <- p0[1] +  DeltaP0
  two ~ dconstraint(p0[2] > 0 & p0[2] < 1)

  alpha2[1] ~ dunif(-1, 5)
  DeltaAlpha ~ dunif(-5, 0)
  alpha2[2] <- alpha2[1] + DeltaAlpha
  one ~ dconstraint(alpha2[2] > -5 & alpha2[2] < 5) # be sure that alpha2[2] is not outside cached arra

  AlphaID[1] <- trunc(alpha2[1]*roundValue+indexValue) # formula (round(alpha2,1)*10+51)  to find were
  AlphaID[2] <- trunc(alpha2[2]*roundValue+indexValue)

  distSq[1:cells, 1:ntraps,1] <- AccessCachedNim(AlphaID[1]) #cached[1:cells, 1:ntraps,AlphaID[1]]
```

```
  distSq[1:cells, 1:ntraps,2] <- AccessCachedNim(AlphaID[2]) #cached[1:cells, 1:ntraps,AlphaID[2]]

  alpha1[1] <- 1/(2 * sigma[1] * sigma[1])
  alpha1[2] <- 1/(2 * sigma[2] * sigma[2])

  ## loop over individuals
  for(i in 1:M) {
    ## AC coordinates
    sxy[i,1] ~ dunif(0, x.max)
    sxy[i,2] ~ dunif(0, y.max)

    sID[i] <- habitatgrid[trunc(sxy[i,1]) + 1, trunc(sxy[i,2]) + 1] # in which cell fall the AC
    eta[i] ~  dbern(probB)
    probcap[i,1:ntraps] <- p0[eta[i]+1] * exp(-alpha1[eta[i]+1] * distSq[sID[i],1:ntraps,eta[i]+1]) # d

    z[i] ~ dbern(psi) # latent dead/alive indicators

    for(j in 1:ntraps){
      muy[i,j] <-  probcap[i,j] * z[i]
      for(k in 1:K){
        y[i,j,k] ~  dbern(muy[i,j])}
    }
  }

  ## derived quantity: total population size
  N <- sum(z[1:M])
})
```

Compute the centroid of detection for each individual detected so his activity center is near it's detection

```
Sxy_inits <- tibble(X = rep(NA,M), Y = rep(NA,M))

# Centroid for detected individuals
for(i in 1:N.det){
  Tl <- traplocs[which(apply(yy[i,,],1, sum) >0),] # trap location were an individual was detected
  if (length(Tl)>2){
    Sxy_inits[i,"X"] <- apply(Tl,2,mean)["X"]
    Sxy_inits[i, "Y"] <- apply(Tl,2,mean)["Y"]} # centroid
  else {
    Sxy_inits[i,"X"] <- Tl["X"]
    Sxy_inits[i, "Y"] <- Tl["Y"]
  }
}

# random values for AC of undetected individuals

for(i in N.det:M){
    Sxy_inits[i,"X"] <- runif(1,0,x.max)
    Sxy_inits[i, "Y"] <- runif(1,0,y.max)
}
```

Define initial values

```
# True values
# nimInits <- list(p0 = c(p0.a,p0.b),
#                  psi = N/M,
#                  sigma = c(sigma.a,sigma.b),
#                  alpha2 = c(alpha2.a,alpha2.b),
#                  probB  = c(N.b/(N.a+N.b)),
#                  DeltaAlpha = DeltaAlpha,
#                  DeltaSigma = DeltaSigma,
#                  DeltaP0 = DeltaP0,
#                  eta = c(rep(NA,N.det), rbinom(M-N.det, 1, prob=0.5)),
#                  sxy = Sxy_inits %>% as.matrix(),
#                  z = c(rep(1,N.det),rbinom(M-N.det, 1, prob=0.3)))

# Random values
p0Inits <- runif(1,0.1,0.4)
alphaInits <- runif(1, 1, 5)
DeltaAlphaInits <- runif(1, -1, -0.1)
sigmaInits <- runif(1, 1, 3)
DeltaSigmaInits <- runif(1, 0.5, 2)
DeltaP0Inits <- runif(1,-0.1,0)
psiInits <- runif(1,0.7,0.9)

nimInits <- list(p0 = c(p0Inits,p0Inits+DeltaP0),
                 psi = psiInits,
                 sigma = c(sigmaInits,sigmaInits+DeltaSigmaInits),
                 alpha2 = c(0,0),
                 probB  = runif(1,0.05,0.2),
                 DeltaAlpha = 0,
                 DeltaSigma = DeltaSigmaInits,
                 DeltaP0 = DeltaP0,
                 eta = c(rep(NA,N.det), rbinom(M-N.det, 1, prob=0.5)),
                 sxy = Sxy_inits %>% as.matrix(),
                 z = c(rep(1,N.det),rbinom(M-N.det, 1, prob=0.3)))
```

Define constants and data to run the model

```
nimConstants <- list(M = M,
                     ntraps = Array*Array,
                     K = K,
                     x.max = x.max,
                     y.max = y.max,
                     cells = nrow(spatialdomain))

nimData <- list(y = yy,
                habitatgrid = habitatgrid,
                eta = eta_data,
                indexValue = indexValue,
                roundValue = roundValue,
                #cached = cached,
                one = 1,
                two = 1,
                three = 1)
```

```r
model <- nimbleModel(code = modelCode2groups,
                     constants = nimConstants,
                     data = nimData,
                     inits = nimInits,
                     check = F,
                     calculate = F,
                     dimensions = list(habitatgrid = dim(habitatgrid),
                                       probcap = c(M, ntraps),
                                       sID = M,
                                       eta = M,
                                       AlphaID = 2))
```

```
## Defining model

## Building model

## Setting data and initial values

## Checking model sizes and dimensions

##   [Note] This model is not fully initialized. This is not an error.
##          To see which variables are not initialized, use model$initializeInfo().
##          For more information on model initialization, see help(modelInitialization).
```

```r
# model$calculate()
# cmodel <- compileNimble(model)
# cmodel$calculate()
# MCMCconf <- configureMCMC(model = model,
#                           monitors = c("N", "sigma", "p0","psi","alpha2","probB"),
#                           control = list(reflective = TRUE),
#                           thin = 1)
#
# MCMC <- buildMCMC(MCMCconf)
# cMCMC <- compileNimble(MCMC, project = model, resetFunctions = TRUE)
# # RUN THE MCMC
# MCMCRuntime <- system.time(myNimbleOutput <- runMCMC(mcmc = cMCMC,
#                                                      nburnin = 1000,
#                                                      niter = 50000,
#                                                      nchains = 2,
#                                                      samplesAsCodaMCMC = TRUE))
#
# save(myNimbleOutput, file = "myNimbleOutput_appendixS1.RData")
load("myNimbleOutput_appendixS1.RData")
```

```r
basicMCMCplots::chainsPlot(myNimbleOutput,var=c("N","p0","sigma","alpha2","psi","probB"), line = c(N,p0
```

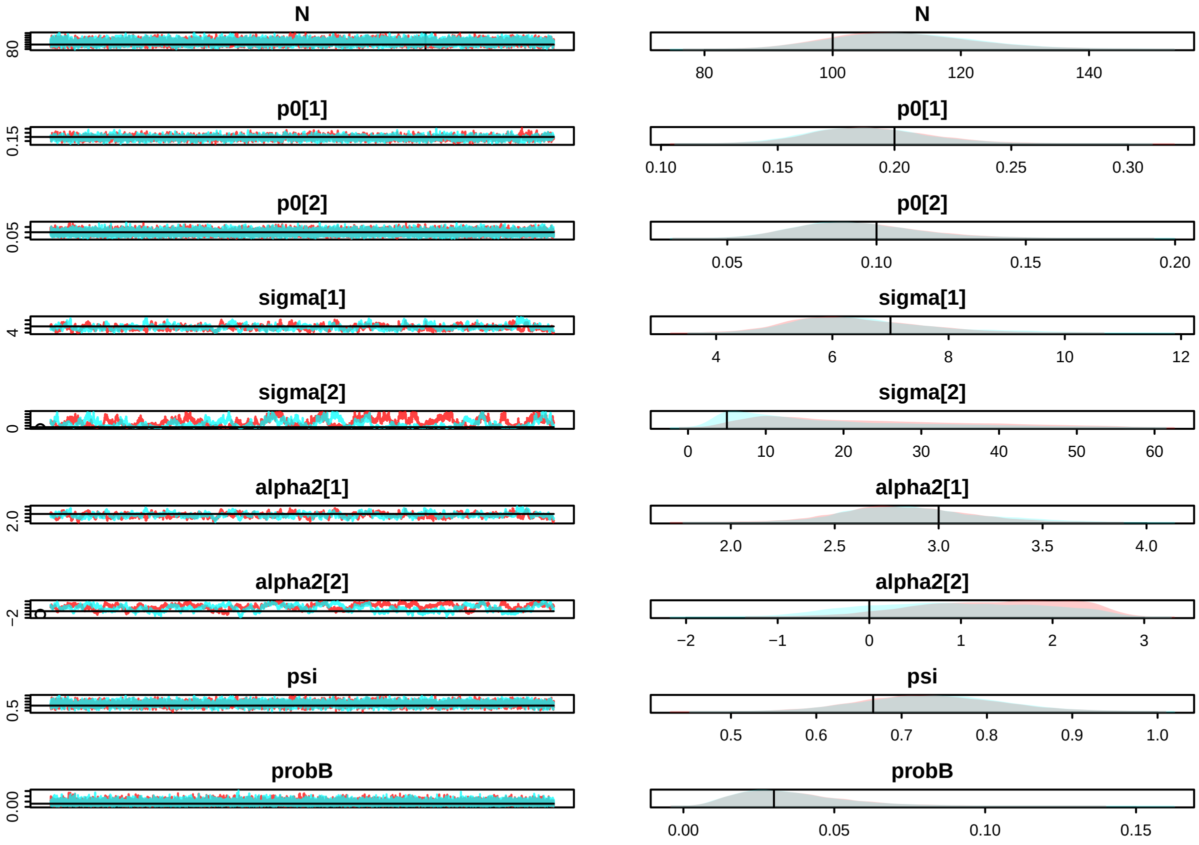


```r
summary(myNimbleOutput)
```

```
##
## Iterations = 1:49000
## Thinning interval = 1
## Number of chains = 2
## Sample size per chain = 49000
##
## 1. Empirical mean and standard deviation for each variable,
##    plus standard error of the mean:
##
##                 Mean       SD  Naive SE Time-series SE
## N          110.84580 10.51847 3.360e-02      0.2035925
## alpha2[1]    2.85215  0.29718 9.493e-04      0.0189300
## alpha2[2]    1.15945  0.89956 2.874e-03      0.0907727
## p0[1]        0.18902  0.02425 7.745e-05      0.0005747
## p0[2]        0.09353  0.01984 6.339e-05      0.0002192
## probB        0.03567  0.01765 5.639e-05      0.0001381
## psi          0.73578  0.07765 2.480e-04      0.0014527
## sigma[1]     6.45123  1.13536 3.627e-03      0.0693649
## sigma[2]    19.67454 12.93414 4.132e-02      1.1735708
##
## 2. Quantiles for each variable:
##
##                 2.5%       25%       50%      75%     97.5%
```

```
## N          92.000000 103.00000 110.00000 118.0000 133.00000
## alpha2[1]  2.314221   2.64611   2.83128   3.0400   3.48523
## alpha2[2] -0.612783   0.50321   1.20869   1.8880   2.60534
## p0[1]      0.145310   0.17193   0.18762   0.2045   0.23980
## p0[2]      0.059751   0.07945   0.09167   0.1058   0.13745
## probB      0.009919   0.02276   0.03276   0.0454   0.07781
## psi        0.588951   0.68161   0.73380   0.7884   0.89239
## sigma[1]   4.595920   5.63898   6.30144   7.1189   9.10922
## sigma[2]   3.767919   9.21338  16.08018  28.0876  49.84539
```

```r
MCMCvis::MCMCsummary(myNimbleOutput)
```

```
##                  mean          sd         2.5%          50%        97.5% Rhat
## N          110.84579592 10.51846981 92.000000000 110.00000000 133.00000000 1.00
## alpha2[1]    2.85214529  0.29717879  2.314221186   2.83128309   3.48522553 1.02
## alpha2[2]    1.15944955  0.89956153 -0.612782924   1.20868799   2.60533518 1.34
## p0[1]        0.18901802  0.02424580  0.145309536   0.18762341   0.23980142 1.01
## p0[2]        0.09352839  0.01984293  0.059751294   0.09167085   0.13744900 1.00
## probB        0.03566711  0.01765322  0.009919494   0.03276369   0.07780762 1.00
## psi          0.73578412  0.07764746  0.588950511   0.73379588   0.89239072 1.00
## sigma[1]     6.45122737  1.13536372  4.595919887   6.30144010   9.10922163 1.04
## sigma[2]    19.67453885 12.93413811  3.767919205  16.08017869  49.84539012 1.24
##            n.eff
## N           2666
## alpha2[1]    252
## alpha2[2]     95
## p0[1]       1776
## p0[2]       8197
## probB      16378
## psi         2858
## sigma[1]     274
## sigma[2]     116
```

#### 2.2 Mixture Model

```r
# # define parameters
# nimConstants <- list(M = M,
#                      ntraps = ntraps,
#                      K = K,
#                      x.max = x.max,
#                      y.max = y.max,
#                      cells = nrow(spatialdomain))
#
# nimData <- list(y = yy,
#                 habitatgrid = habitatgrid,
#                 indexValue = indexValue,
#                 roundValue = roundValue,
#                 cached = cached,
#                 one = 1,
#                 two = 1)
#
```

```
# probBInits <- runif(1,0.05,0.2)
# nimInits <- list(p0 = c(p0Inits,p0Inits+DeltaP0),
#                  psi = psiInits,
#                  sigma = c(sigmaInits,sigmaInits+DeltaSigmaInits),
#                  alpha2 = c(0,0),
#                  probB  = probBInits,
#                  DeltaAlpha = 0,
#                  DeltaSigma = DeltaSigmaInits,
#                  DeltaP0 = DeltaP0,
#                  eta = rbinom(M, 1, prob=probBInits),
#                  sxy = Sxy_inits %>% as.matrix(),
#                  z = c(rep(1,N.det),rbinom(M-N.det, 1, prob=0.3)))
#
# model <- nimbleModel(code = modelCode2groups,
#                      constants = nimConstants,
#                      data = nimData,
#                      inits = nimInits,
#                      check = F,
#                      calculate = T,
#                      dimensions = list(habitatgrid = dim(habitatgrid),
#                                        probcap = c(M, ntraps),
#                                        sID = M,
#                                        AlphaID = 2))
```

```
# calculate(model)
# cmodel <- compileNimble(model)
# MCMCconf <- configureMCMC(model = model,
#                           monitors = c("N", "sigma", "p0","psi","alpha2", "probB"),
#                           control = list(reflective = TRUE),
#                           thin = 1)
#
# MCMC <- buildMCMC(MCMCconf)
# cMCMC <- compileNimble(MCMC, project = model, resetFunctions = TRUE)
#
# # RUN THE MCMC
# myNimbleOutput <- runMCMC(mcmc = cMCMC,
#                           nburnin = 1000,
#                           niter = 10000,
#                           nchains = 2,
#                           samplesAsCodaMCMC = TRUE)
```

```
# basicMCMCplots::chainsPlot(myNimbleOutput,var=c("N","p0","sigma","alpha2","psi","probB"), line = c(N,
#
# summary(myNimbleOutput)
```
