## Appendix S2 for "Unravelling the effects of heterogeneity in space use on estimates of connectivity and population size: Insights from spatial capture-recapture modelling"

### Simulation Study

***Table S2.A****: Table with prior distribution for each model parameter*

| Parameter | Distribution |
|---|---|
| sigma[1] | Uniform(0, 50) |
| DeltaSigma | Uniform(-50, 50) or Uniform(-50,150) for model MO3 |
| sigma[2] | sigma[1]+DeltaSigma |
| alpha2[1] | Uniform(-1, 5) |
| DeltaAlpha | Uniform(-5, 0) |
| alpha2[2] | alpha2[1]+DeltaAlpha |
| p0[1] | Uniform(0,1) |
| DeltaP0 | Uniform(-1,0) |
| p0[2] | P0[1]+DeltaP0 |
| psi | Uniform(0,1) |
