## Appendix S3 for "Unravelling the effects of heterogeneity in space use on estimates of connectivity and population size: Insights from spatial capture-recapture modelling"

<u>Case study</u>:

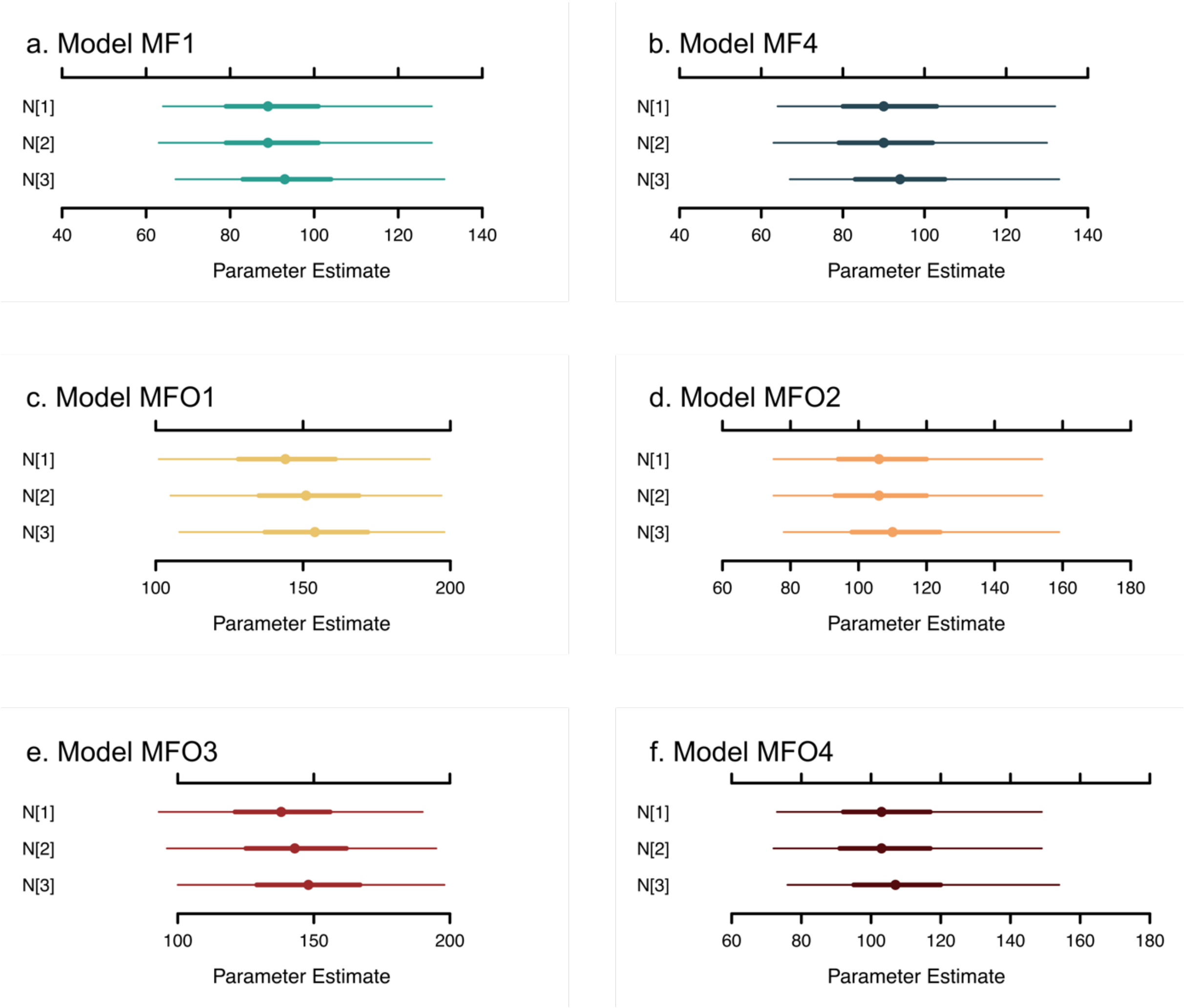


***Figure S3.A**: Estimated population size in 2017 (N[1]), 2018 (N[2]) and 2019 (N[3]) for the Pyrenean brown bear population. Caterpillar plots display the median of the posterior distribution and the 50% and 95% credible intervals for the 6 tested models. The first row shows the models with outliers removed i.e. MF1 (a) and MF4 (b) and the second and the third rows show the models with all the dataset i.e. models MFO1 (c), MFO2 (d), MFO3 (e) and MFO4 (f).*

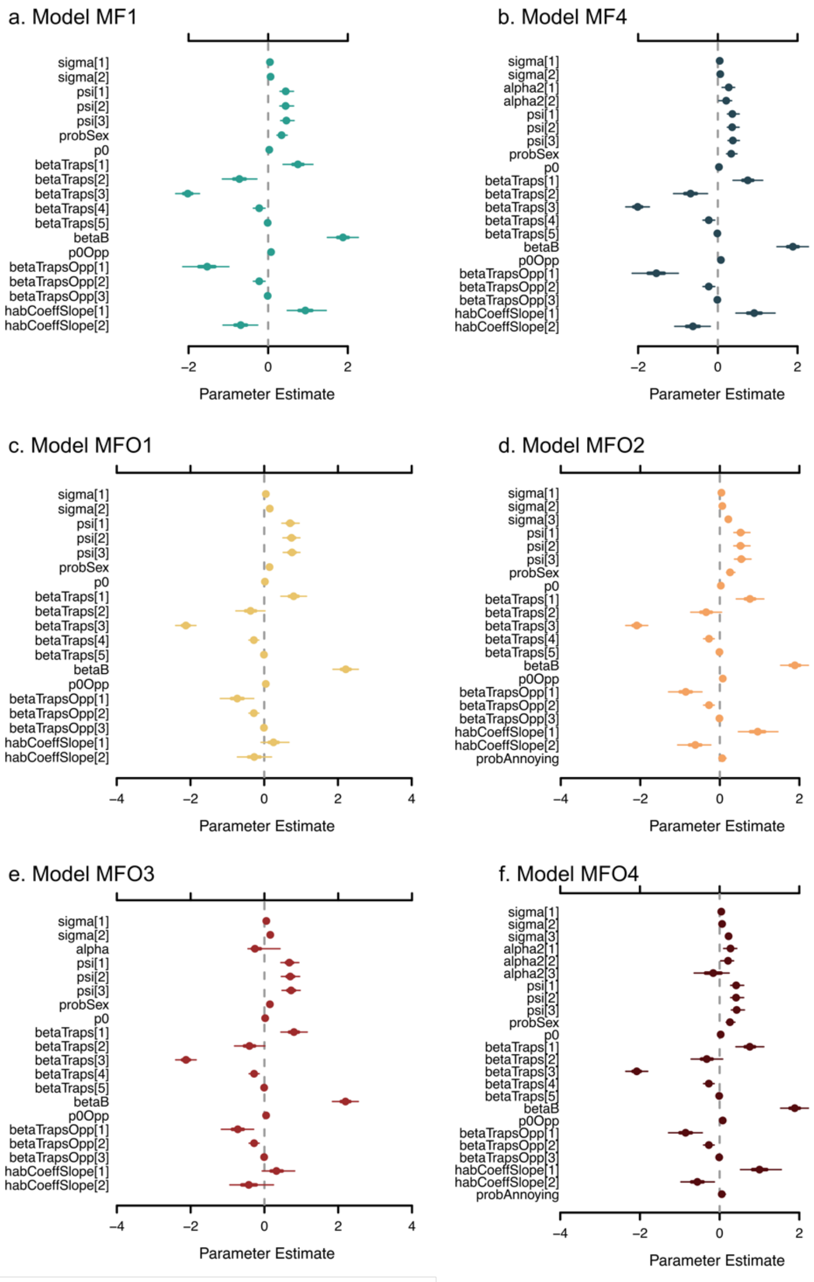


***Figure S3.B**: Estimated parameters for the Pyrenean brown bear population. The parameters are the scale parameter for females (sigma[1]) and males (sigma[2]), the resistance parameters for females (alpha2[1]), males (alpha2[2]) and outliers (alpha2[3]), the inclusion parameters in 2017(psi[1]), 2018 (psi[2]) and 2019 (psi[3]), the probability to be a male (probSex), the intercept of the baseline detection probability for the structured monitoring (p0), the slope for the number of visits (betaTraps[1]), the slope for the country for the structured monitoring (betaTraps[2]), the slope for the type of trap (betaTraps[3]), the slope for effect of the month (betaTraps[4], betaTrapsOpp[2]) and the squared effect of the month (betaTraps[5], betaTrapsOpp[3]), the slope for the behavioral effect (betaB), the intercept of the baseline detection probability for the opportunistic monitoring (p0Opp), the slope for the country for the opportunistic monitoring (betaTrapsOpp[1]), the habitat slope for the ruggedness (habCoeffSlope[1]) and the habitat slope for the human density (habCoeffSlope[2]).*
*Caterpillar plots display the median of the posterior distribution and the 50% and 95% credible intervals for the 6 tested models. The first row shows the models with outliers removed i.e. MF1 (a) and MF2 (b) and the second and the third rows show the models with all the dataset i.e. models MFO1 (c), MFO2 (d), MFO3 (e) and MFO4 (f).*

***Table S3.A**: Table with prior distribution for each model parameter*

| Parameter | Distribution |
|---|---|
| sigma | Uniform(0, 50) |
| alpha2[1] | Uniform(-1.5, 1.5) |
| DeltaAlpha | Uniform(-1.5, 0) |
| alpha2[2] | alpha2[1]+DeltaAlpha |
| DeltaAlpha2 | Uniform(-1.5, 0) |
| alpha2[3] | alpha2[2]+DeltaAlpha2 |
| p0 | Uniform(0,1) |
| betaTraps | Uniform(-5,5) |
| betaB | Uniform(-5,5) |
| p0Opp | Uniform(0,1) |
| betaTrapsOpp | Uniform(-5,5) |
| habCoeffSlope | Normal(0,10) |
| probSex | Uniform(0,1) |
| probOutlier | Uniform(0,1) |
